# Indexing and searching petabyte-scale nucleotide resources

**DOI:** 10.1101/2023.07.09.547343

**Authors:** Sergey A. Shiryev, Richa Agarwala

**Affiliations:** Department of Health and Human Services, National Center for Biotechnology Information, National Library of Medicine, National Institutes of Health, Bethesda MD 20894

## Abstract

Searching vast and rapidly growing sets of nucleotide content in data resources, such as runs in Sequence Read Archive and assemblies for whole genome shotgun sequencing projects in GenBank, is currently impractical in any reasonable amount of time or resources available to most researchers. We present Pebblescout, a tool that navigates such content by providing indexing and search capabilities. Indexing uses dense sampling of the sequences in the resource. Search finds subjects that have short sequence matches to a user query with well-defined guarantees. Reported subjects are ranked using a score that considers the informativeness of the matches. Six databases that index over 3.5 petabases were created and used to illustrate the functionality of Pebblescout. Here we show that Pebblescout provides new research opportunities and a data-driven way for finding relevant subsets of large nucleotide resources for analysis, some of which are missed when relying only on sample metadata or tools using pre-defined reference sequences. For two computationally intensive published studies, we show that Pebblescout rejects a significant number of runs analyzed without changing the conclusions of these studies and finds additional relevant runs. A pilot web service for interactively searching the six databases is freely available at https://pebblescout.ncbi.nlm.nih.gov/

## INTRODUCTION

The mission of the National Center for Biotechnology Information (NCBI) is to develop new information technologies to aid in the understanding of fundamental molecular and genetic processes that control health and disease. NCBI is home to many well-known and freely available databases and tools, including PubMed, GenBank, Sequence Read Archive (SRA), and BLAST®. Sequences in GenBank include assemblies submitted for the whole genome sequencing (WGS) projects. NCBI’s Reference Sequence (RefSeq) collection provides a comprehensive, integrated, non-redundant, well-annotated set of sequences, including assemblies. In this manuscript, we use WGS and RefSeq to refer to the set of assemblies from the corresponding projects and collection, respectively.

SRA and WGS are examples of two large nucleotide sequence resources available at NCBI. As of February 2023, SRA had over 40 petabases of publicly available sequence data in over 23.7 million sequencing *runs*, and WGS had over 1.5 million assemblies. Each sequencing run contains many *reads*, usually at least tens of thousands, and a WGS assembly may contain hundreds of *contigs*. Currently, most of the reads in SRA are at most 250 bases long while contigs in WGS are at least 200 bases long and can be a few hundred megabases in length.

SRA and WGS can be resources for discovery, but because of their size, it is impractical to provide public sequence search capabilities for all data in these resources. Currently, the main approaches for analyzing such resources fall in following four categories:

1. Create an index on the resource that is very sparse and/or removes some data^1–4^
2. Find potentially interesting subset of the resource via a proxy for a direct sequence match, such as, by using sample metadata^5, 6^, taxonomy information^5, 7, 8^, or marker genes^9, 10^
3. Arbitrarily limit the number of reads analyzed from a run^11, 12^
4. Accept the high cost of doing analysis but ease the process of running tools on large compute clusters^5, 11^, or develop pipelines that use special hardware^13^

The need for general-purpose solutions accessible to any researcher for analyzing large resources is well known^14^. The BLAST web service available at NCBI is an example of such a solution for searching some large databases; however, search against SRA and WGS using BLAST web service requires users to limit the search to a subset of these resources using some property, such as taxonomy for WGS and experiment for SRA. The ability to find relevant subsets of the resource for mining information is critical (i) for materializing the benefits anticipated when these resources were created and (ii) for justifying the ongoing cost of sequencing and supporting these rapidly growing resources.

We present Pebblescout (**P**ick **E**nough **B**its to **B**uild **L**ong **E**nough **S**equence **C**lues **O**n **U**ser **T**arget), a suite of software tools that can be used for (i) indexing sequence data in a resource *once* and (ii) searching the index to produce a ranked list for the subset of the resource with matches to *any* user query; the guarantee on the match length is determined by the parameters used for indexing. If necessary, a user can then perform additional computations, such as alignment or assembly, on a much smaller subset of the large resource. A user may also find patterns in metadata of the relevant subset reported that may not be detectable otherwise. The four main advantages of Pebblescout compared to current approaches for indexing[1–4] and finding subsets[5–10] are that (i) the index is dense with, on average, information saved from every ninth position of the sequences with parameters we used, (ii) all sequences in the resource are used for indexing, (iii) the search has a guarantee for match length, and (iv) the search points to the subset of the resource using sequence information in the user query. The main resource requirement for Pebblescout is a network attached random access storage array for storing the index.

Indexing uses sampling, encoding, hashing, and compression techniques. Together, the index and metadata for *subjects* are referred to as a *database*. The search component of the system generates hashes from the user query using the same parameters used for creating databases. Subjects are ranked in the output. Ranking reflects informativeness of hashes sampled from the query; the informativeness of a hash is measured by the number of subjects from which the same hash was sampled in the database.

To illustrate the capabilities of Pebblescout, we created six databases: four from publicly available runs in SRA and one each for RefSeq and WGS (Table 1). Four databases from SRA have subjects that are metagenomic runs released before the end of 2021, Human RNAseq runs released in 2021, and the final two are comprised of runs from Phase 3 of the 1000 Genomes Project^15^ indexed differently – the first where each run is a subject and the second where all runs for the same BioSample are considered one subject. The WGS and RefSeq databases contain assemblies available as of Feb 14, 2022 and April 22, 2022, respectively, in the corresponding resources. Subject identifiers in the databases are run identifiers, BioSample identifiers, or assembly accessions depending on the unit considered as one subject. A pilot website where users can submit queries and interactively search these databases is available at https://pebblescout.ncbi.nlm.nih.gov/.

**Table 1:** Pebblescout indexes (Pb: petabases; Tb: terabases; TB terabytes)

| Database | #Subjects | Total Bases (B) | DB size (D) | B / D | Cutoff date |
| --- | --- | --- | --- | --- | --- |
| Metagenomic Runs | 3,236,120 | 1.75 Pb | 108 TB | 16.2 | 12/31/2021 |
| WGS assemblies | 1,144,201 | 17.13 Tb | 5.4 TB | 3.2 | 02/14/2022 |
| RefSeq assemblies | 257,586 | 2.2 Tb | 0.8 TB | 2.7 | 04/22/2022 |
| Phase3 runs | 36,841 | 612 Tb | 28.9 TB | 21.2 | 03/14/2022 |
| Phase3 BioSamples | 2,503 | 612 Tb | 4.2 TB | 145.7 | 03/14/2022 |
| Human RNAseq Runs | 285,417 | 1.13 Pb | 30.8 TB | 36.7 | Released in 2021 only |

With the parameters we chose for creating the databases, usually, the size of the database is at least one order of magnitude smaller than the total length of sequences in SRA runs in the database, but only about a third smaller for assemblies due to relatively low redundancy in assemblies. These databases support search guarantee of no false negatives for 42 base pair (bp) matches. That is, if there is an exact match between a user query and a subject that is at least 42 bases long, that match will be found by the search. Herein, we provide an overview of the indexing process, publicly available databases, search modes, and ten applications that illustrate the impact Pebblescout can have on a variety of problems. Four applications are from published studies that analyzed runs in SRA while the remaining six are examples of potential research applications using a wide range of queries, such as, short sequences with single nucleotide polymorphisms, antimicrobial resistance genes, hard-to-sequence genomes, and microbial communities.

For the two computationally intensive published studies of viral discovery and *Candida auris* detection, we show that Pebblescout reported (i) a significantly smaller subset of metagenomic runs from SRA to analyze than the subset found in the studies using sample metadata and species information, (ii) found all runs in the metagenomic database that the authors found as “good” as per their analysis, and (iii) found additional good runs not reported by the authors. For the other two applications from publications, we show how Pebblescout can be used to enhance those efforts. For the first, analysis using Pebblescout expands knowledge of host species for *Wolbachia* by finding sixteen additional host species, while for the second, analysis using Pebblescout provides context and close matches for ssRNA phage sequences reported as novel in the publication by searching WGS.

These applications and the empirical analysis on false positive rates show that, in practice, using relatively dense indexes as we generated, sampling technique we used, and filtering of Pebblescout output as appropriate for the application leads to very few subjects that are not relevant for the application. It also provides an opportunity to do a more thorough and data-driven analysis of the large nucleotide resources.

### Indexing

Indexing has three main stages:

1. **Harvesting 25-mers**: One hash value is generated for every 25-mer in each subject where the 25-mer or its reverse complement is deterministically chosen for generating the hash. A 25-mer is encrypted using a Feistel cipher, adapted from the method published by Schneier^16^, with the resulting ciphertext used as hash value for the 25-mer. The 25-mer with the smallest hash from all eighteen 25-mers in every 42-mer is saved in the database. Saved 25-mers are called *sampled* from the subject by the *harvesting* process. Only the fact that a 25-mer is sampled from the subject is stored and not the positions nor the number of times the 25-mer is present in the subject.
2. **Aggregating 25-mers**: Subject identifiers are encoded using Huffman code^17^ with the number of distinct 25-mers harvested from a subject as its weight for the encoding. For each stored 25-mer, one merged list of encoded subject identifiers for the subjects from which the 25-mer was harvested is saved.
3. **Creating block structure**: A two-strata B+ tree indexing structure is created on aggregated 25-mers to reduce the number of relatively slow block access operations during search. The top of the tree is stored locally, while leaves reside in a network attached random access storage array.

Additional technical details are available in the Methods section.

Comparison of the two 1000 Genomes Phase 3 databases demonstrates that the size of the database when indexing by sample (as defined by BioSample) is significantly smaller compared to indexing by runs, as the bulk of the space occupied by an index is used for storing the list of subjects for 25-mers. In addition to saving space, creating databases that consider runs from the same BioSample, or very similar assemblies, as one subject may have additional analytical advantages for some applications. Supplementary material provides additional information for the list of subjects indexed in each database (Section: Datasets and Metadata, Table S1). Supplementary material also illustrates some properties of the indexing method using data from the indexes created, and practical considerations from our experience in building the databases (Section: Observations from the six databases).

### Searching

Pebblescout supports three search modes: Profile, Summary, and Detailed. The search in all modes samples 25-mers from the query using the same harvesting process as in indexing. We use kmer to mean 25-mer in the rest of the article, unless specified otherwise.

A *kmer match* means that the kmer sampled from the query was also sampled from the subject. Since the same kmer is sampled from a 42-mer and its reverse complement, and the same harvesting process is used on both query and subject sequences, if there is a 42 bp match between a query and a subject, it will result in a kmer match.

Some queries generate an extremely large number of matches in some databases, such as searching the *E. coli* genome against the metagenomic database. For some other applications, it may be useful to consider only the most informative kmers found in a limited number of subjects. In such instances, it is useful to limit the search in Summary and Detailed modes by *masking* (ignoring) kmers that are sampled from more subjects than the specified count threshold. A default count per database is provided but is also user configurable.

**Profile search** shows the number of subjects with a match in the database for each kmer sampled from the query, providing a high-level view for common and rare parts of the user query.

**Summary search** scores all subjects that match at least one unmasked kmer sampled from the query. Each kmer present in a database is assigned a kmer score that decreases as the number of subjects in the database matching the kmer increase; the rate of decrease is controlled by a user configurable scoring constant. Pebblescout score considers only unmasked kmers sampled from the query. The score for a subject normalizes the sum of kmer scores for all kmers considered from the query that match the subject. The score is normalized (i) by the total kmer score for all kmers considered from the query and (ii) by the fraction of kmers considered from the query that match any subject. A *perfect* Pebblescout score of 100 means that all kmers considered from the query were also harvested from the subject. The methods section provides additional details for the score computation. Output from summary search ranks the subject by Pebblescout score in descending order for each query.

**Detailed search** prints one line for every combination of query, subject, kmer where the kmer considered from the query has a match to a subject. It also prints one line for all kmers sampled from the query that are masked and one line for all kmers that result in no match to any subject in the database. The pilot website requires users to specify a set of subjects explicitly; it reports matches to only the specified subjects with no change in how sampled kmers that are masked or have no match are reported. All lines for each query-subject pair matches are consecutive in the output and sorted by the kmer position in the query.

Output formats for search results are described in the Supplementary material (Sections: Output format for Profile search, Table S2; Output format for Summary search, Table S3; Output format for Detailed search, Table S4).

### Search Applications

We present an overview for four applications from publications that analyzed runs in SRA and six research applications to demonstrate the breadth of Pebblescout’s utility in analyses of large datasets. The accompanying supplementary material provides an in-depth analysis for these applications (Section: Analysis for search applications, Figures S1 and S2, Tables S5 to S9), a discussion on false positives (Section: False positive rate in results with a perfect Pebblescout score), and guidelines for using the pilot web service along with information for performing searches using the web service for all applications presented in the manuscript (Section: Pilot web service and searches for applications, Figure S3, Table S10).

### Find previously undiscovered species related to known species (viral discovery example)

To our knowledge, the largest petabyte-scale sequence alignment to SRA was done for viral discovery using the Serratus pipeline^5^. In this analysis, the nucleotide search component of the pipeline aligned 12,295 queries to over 3.8 million runs. The list of runs was generated using metadata from metagenomic, genomic, and RNAseq runs, and by using species information available in SRA using STAT^7^. The total number of runs assembled using coronaSPAdes^18^ was 56,511 of which 14,304 runs came from the nucleotide search. The authors deemed assemblies for 11,120 runs as “good” for further analysis.

Our analysis with Pebblescout using the same 12,295 queries against the metagenomic index shows the following:

1. Pebblescout summary search filters out >84% of the runs present in the metagenomic index and assessed in the publication. The subset of 3.8 million runs present in the metagenomic index is 1.6 million, of which 1.39 million runs are filtered out by the Pebblescout search. The search also filters out >86% of the metagenomic runs not assessed in the publication (1.36 million out of 1.58 million runs).
2. All 1,815 runs present in both the metagenomic index and the set of 14,304 runs reported by the published nucleotide search are found by Pebblescout. That is, using Pebblescout search to filter metagenomic runs would not have caused any loss of viral discovery reported in the study.
3. Pebblescout finds 1,206 out of 1,207 runs present in both the metagenomic index and the set of 11,120 good assemblies reported. The assembly reported by the authors of [5] for the one run not found by Pebblescout has only one contig. This contig has a strong match to a bacterial species and is unlikely to be a viral sequence.
4. Some runs found by Pebblescout not included by the authors of [5] in their set of good assemblies, when assembled using coronaSPAdes, produce contigs that appear to be novel coronavirus sequences.

Hence, we conclude that Pebblescout reported all good runs identified in the viral discovery study that were present in the Pebblescout metagenomic index, potentially identified several additional runs of value, and did so while substantially rejecting many runs that were aligned in one step of the Serratus pipeline.

### Find known species in new environments (*Candida auris* example)

A recent study^6^ aligned the ITS region of *C. auris* as query to 603,692 runs in an attempt to identify environmental niches of *Candida auris*. The runs were identified using keywords “fungi, ITS1, ITS2, mycobiome”. The analysis found *C. auris* in 34 runs. 19 of these 34 runs are in the Pebblescout metagenomic database; the remaining 15 runs are excluded because they are tagged as “genomic” in the SRA metadata.

To find a match count threshold that separates species-specific kmers from other species, we did a Pebblescout profile search to the metagenomic index using the same ITS region as query and identified 76 as the appropriate masking threshold. Using this threshold, a Pebblescout summary search finds 129 runs, including all 19 runs identified in the published study present in the metagenomic database. Analysis using read alignments for the 129 runs found by Pebblescout shows that 86 of them have full-length alignments to species-specific regions of ITS, including 7 runs from three BioProjects not reported by the authors of [6] that are likely of high interest as environmental niches for *C. auris*.

Hence, we conclude that use of Pebblescout for this study, had it been available at the time, would not only have identified the runs reported from the metagenomic database, but would have reduced the computational effort required, and would have returned additional runs of interest.

### Find host species for endosymbionts (*Wolbachia* example)

A recent study^12^ used 44 reference transcripts for three genes (*ftsZ, groE, wsp*) from 36 host species to identify additional arthropod host species for *Wolbachia*. The authors found 2,545 runs from 288 host species in SRA using various metadata criteria available in “Run Selector” provided on the SRA web page. They also used an iterative bait and assemble strategy for assembling genomes using seven reference genomes representing *Wolbachia* isolates from insects and nematodes.

A Pebblescout summary search was done using the same 44 reference transcripts as queries to the metagenomic database. The analysis of the search results presented in the supplementary material found 16 host species not found by the authors of [12]. Of the 16 host species, 11 host species contained all three genes, 4 host species contained 2, and 1 host species contained only 1. For each of the host species identified, the run whose assembly produced the maximum coverage to a reference genome from all runs for the same species had reference genome coverage in the range from 1% to 90%.

This analysis demonstrates that Pebblescout can be used to create an initial list of subjects based on sequence information that can be subsequently filtered using the study criteria. Doing so the other way around, as was done in the published study, leads to missing some interesting runs and an incomplete analysis of the data publicly available in SRA.

### Find assemblies in WGS for additional context or to support novel sequence findings (ssRNA phage example)

A study^19^ to expand the set of known ssRNA phage genomes identified 15,611 novel sequences from 82 publicly available metatranscriptomic datasets generated from activated sludge and aquatic environments. Using these 15,611 sequences as queries, Pebblescout finds one WGS assembly to which hundreds of queries have a full-length alignment with ≥95% identity. This example shows that Pebblescout provides an easy and effective way to find assemblies in WGS that have contigs with similar sequence as the given query. Such an analysis can provide additional context for the new sequences or show that nothing similar exists in WGS.

### Research applications from a wide range of queries

We present a brief overview for six different research problems that can utilize Pebblescout. Details for these applications are available in the supplementary material.

1. **Detect hard-to-sequence samples (*Cyclospora cayetanensis* example):** We detect partial sequences for a hard-to-sequence genome, *Cyclospora cayetanensis*, in SRA by doing Detailed search on the metagenomic index using 5,793 RefSeq mRNA sequences annotated on the *C. cayetanensis* assembly as queries. We show that by removing spurious matches using information in the detailed search, we find 165 transcript-Run pairs where at least five reads from the runs have good alignments to the transcript. This example shows that Pebblescout may prove to be a particularly valuable tool for the identification of reservoirs containing limited sequences of genomes, such as those associated with difficult-to-sample species.
2. **Find variant/invariant regions of genomes (*E. coli* and HIV-1 examples):** We find common and rare parts of *E. coli K12* and HIV-1 genomes by doing a Profile search using genomes queried against metagenomic and Human RNAseq databases, respectively. The *E. coli* profile has five sampled positions with no more than 250 matches. These positions, seen rarely in the metagenomic database, may be useful in applications, such as pathogen detection and understanding evolution of strains over time. The HIV-1 profile analysis shows high-frequency positions that are likely due to cloning vectors and positions with higher variability; one of these is gp160 region that is known to be variable.
3. **Find contamination (SARS-CoV-2 example):** We show that Pebblescout can be used to find contaminating sequences from other taxa in WGS by using SARS-CoV-2 genome as the query. Our analysis finds 36 *E. coli* assemblies in WGS from the same sequencing project that have contigs matching SARS-CoV-2 genome. Removing contamination is essential to avoid making incorrect judgements using assemblies.
4. **Find subjects with genes of interest (Methicillin-resistance genes example):** We assess the presence of methicillin-resistance genes in RefSeq and WGS assemblies and metagenomic runs using complete coding sequences for mecA, mecB, mecC, and mecD genes as queries. Analysis shows that for each of the 1,076 query-assembly pairs to RefSeq and 1,983 query-assembly pairs to WGS with a perfect Pebblescout score, there are alignments between the query transcript to one or more contigs in the assembly that are at 100% identity for the entire length of the query. Assessment of 1,422 query-Run pairs to metagenomic index with perfect Pebblescout score also shows that Pebblescout identifies excellent candidates for use in assembling these antimicrobial resistance genes.
5. **Find subjects with SNPs of interest (*Acinetobacter baumannii* example):** We generated queries using six single nucleotide polymorphisms (SNPs) reported for bacterial pathogen *Acinetobacter baumannii* that differentiate two published outbreak clusters. The queries were of length 201 bp, 83 bp, and 49 bp with the polymorphic base as the middle base. Analysis of subjects reported with perfect Pebblescout score using the variant base in queries against RefSeq, WGS, and metagenomic databases shows that 100% of RefSeq assemblies, 74 out of 75 WGS assemblies, and 244 out of 254 metagenomic runs support the variant base. There were no incorrect subjects reported with queries of length 201 bp, six metagenomic runs with queries of length 83 bp, and one WGS assembly and 10 metagenomic runs with queries of length 49 bp. This analysis shows we can find subjects with SNPs of interest by creating queries using the SNP base and that the false positive rate is low in the matches reported with perfect Pebblescout score.
6. **Find similar large and heterogeneous samples (microbial community example):** To find microbial communities similar to those in a water sample from the Chesapeake Bay, we randomly selected one hundred thousand reads from SRR14874039 and concatenated them with ambiguity character ‘N’ to generate one query. Searching the metagenomic database with this query and categorizing location for top 50 scoring samples shows that all except one sample are geographically close to Chesapeake Bay of which 30 are from the Chesapeake Bay itself. To make the search more specific to Chesapeake Bay, we show that by changing score constant to 1 (essentially ignoring all but very rarely seen kmers), locations for top 50 scoring samples include 45 samples from the Chesapeake Bay. This analysis suggests that Pebblescout identifies runs with a microbial community similar to the one in the input query and allows for different questions to be answered for the same input with different parameters, when the size and properties of the input support finding such differences.

## Discussion

We present Pebblescout as a new tool that can index large nucleotide resources and search those databases. Search produces a ranked list of the content in the resource with matches to the user query. The search can be customized based on the application to be very specific or broad. Three levels of search results are available to give users a high-level overview, a summary, or low-level details for the results. Six databases were created to illustrate Pebblescout’s functionality that indexed over 3.5 petabases of sequence data. These databases can be interactively searched free of cost by any researcher with rapid turnaround by submitting queries to the Pebblescout pilot website https://pebblescout.ncbi.nlm.nih.gov/.

We demonstrate through re-analysis of several publications that Pebblescout is an effective tool that can reduce the volume of sequences used for the corresponding analyses without changing the conclusions of these studies, thereby reducing the time and resources spent without sacrificing quality. We also show that Pebblescout provides a data-driven way to find relevant subsets of content in vast and rapidly growing sequence databases. Doing so enhances the quality of results in some studies while in other cases, an evaluation of the sample metadata available for search results is informative. It may also spur additional studies.

As the first general purpose solution supporting agile navigation of petabyte-scale nucleotide sequence sets, Pebblescout becomes a benchmark for future solutions to this problem. Future work may include improvements to scoring methods, presentation of outputs, and integration with other analysis tools. We encourage users to provide feedback in the context of their use cases on the Pebblescout website or by sending email to.The information will inform future development of this tool.

## METHODS

### Database building and index architecture

The three main stages in building an index are sampling 25-mers from each subject (called *harvesting*), aggregating 25-mers across all subjects, and creating a block structure for navigating the index. Additional steps in building a database for verifying harvested 25-mers, incorporating metadata, determining default values for score constant and masking threshold, and for updating a database are also described below.

Supplementary information provides information for various steps from the databases we built, such as evidence for the even distribution of kmers, memory needed for harvesting, importance of verification, and choice of alphabet (Section: Observations from the six databases).

### Harvesting 25-mers

The two-bit encoding used for the four DNA letters A, C, G, T is 00, 01, 10, 11, respectively. For every 25-mer with no ambiguous bases, either the 25-mer or its reverse complement is chosen so that the middle base is always an A or a G. That is, if the middle base of the 25-mer is T or C, then its reverse complement is chosen so that the middle base becomes A or G, respectively. With this choice, the right bit for the 13^th^ base in the 50 bits for the 25-mer is always a zero. This bit is discarded and the remaining 49 bits (the left 25 bits and the right 24 bits of the 50 bits) are used for Feistel encoding. This creates a pseudo-random 1-to-1 mapping for the 49 bits where the mapping has no collisions (a perfect hash function) and is also inversible.

We use the rightmost seven bits of the 49-bits to divide the encodings into 128 disjoint sets, referred to as 128 *slices* of the index. Using the printable alphabet of size 64, the 42 remaining bits are translated to a string of size 7 as each character in the alphabet codes for 6 bits. The alphabet contains ten base 10 numerals, 26 lowercase and 26 uppercase letters in the English alphabet, and two curly brackets. The rightmost seven bits from the Feistel encoding distribute 25-mers evenly in the 128 slices without *a priori* knowledge of the distribution of 25-mers in the subjects. Even distribution of 25-mers in slices is important for indexing and search efficiency, as it allows for good load balancing and for the parallel processing of each slice to take approximately the same time and space.

From every 18 consecutive 25-mers, the 25-mer with the smallest ciphertext *(minimizer)* is saved in the index. That is, from every 42-mer without any ambiguities, at least one 25-mer is saved in the index. The same 25-mer can be chosen by more than one 42-mer. Additionally, more than one 25-mer from a 42-mer can appear in the index, as another 42-mer that overlaps it by at least 25 bases can choose a different 25-mer from the overlapping portion. The index does not store positions nor the number of times a 25-mer was harvested from the subject. Omitting positions is necessary to get compression.

Each row in the index file for each slice has the first 7 characters for the 25-mer as described above. These rows are sorted lexicographically in the output. To process the data efficiently, we combine the collection of minimizers with sorting them. For the 49-bit words, using MSD Radix on a Judy array in a bitmap form to sort takes linear time compared to the typical O(nlog(n)). Furthermore, the Judy array’s memory requirement is modest. This is because minimizers not only reduce the amount of data to be stored, but more critically, they skew the distribution of 49-bit words towards small integer values; the skew is exploited by Judy array to use memory efficiently. The distribution with counts of kmers binned by prefixes of sizes one, two, and three show a steep decay that makes the top level of the radix tree very sparse.

As a check, a row at the end of the index file for each slice is added with 7 tildes (called *endcap*). Tilde is not part of the 64 characters and is lexicographically larger than any of the characters in the alphabet of 64. This endcap is used as an indicator of a successful execution. A second file that gives the number of rows in the file, excluding the row for endcap, is also produced.

### Verifying 25-mers

When harvesting of 25-mers is done on many subjects, it requires reading the sequence for those subjects over the network and writing two orders of magnitude more files over the network. In such a situation, hardware and network glitches not caught by the computing infrastructure are possible. To reduce the chance of storing wrong information in the index, we re-run the harvesting step a second time using an option that only produces the counts instead of both counts and the encoding for 25-mers. If any of the 128 counts for any subject do not match between the two executions, that subject is harvested and verified again. We also check for the presence of the endcap in the last row. These additional checks are necessary to avoid producing syntactically correct files that contain incorrect information.

### Aggregating 25-mers

Each slice is processed independently. For each slice and subject in the database, we know the number of 25-mers saved from the subject in the slice. Subject identifiers are encoded using Huffman code with the number of 25-mers harvested from the subject in the slice as its weight for the encoding. Note that the same subject may have a different encoding in different slices.

For each stored 25-mer, one merged subject list for all subjects the 25-mer was sampled from is saved using the respective encodings. Compression is achieved as short codes are assigned to subjects that have many 25-mers in the index, and long codes to subjects with only a few 25-mers. The string of encoded subject identifiers is decoded from left to right without a need for any delimiter separating the encodings.

The alphabet used for *n*-ary Huffman coding can be ASCII printable characters or binary. For printable alphabet, we use *n* of 92 printable characters that excludes backslash, double quotes, delete, and space from the 96 printable characters. For binary alphabet, we use *n* of 252 characters excluding null, carriage return, line feed, and space. In both cases we use byte-sized literals.

### Creating block structure

The files with aggregated 25-mer lists have rows stored in a lexicographically sorted order. The entire file is viewed as a sequence of blocks not exceeding 4,096 bytes in size, except for the rows that have subject lists exceeding 4,096 bytes. In those cases, associated blocks contain only one row and are as big as necessary. To further reduce the *main index* created from the aggregated 25-mer files, an additional requirement was added: for all rows within each block, the 4-character prefix must be the same and thus can be dropped. That is, for each row of the aggregated 25-mers, the suffix starting from the fifth character is saved in the main index and order of rows maintained as it was in the sorted aggregated list. For example, 19nbIcH}t#}uJ}v4 appears on row 420,904,771 of the aggregated 25-mers in the first slice of the WGS database. Prefix 19nb is used in the creation of the block structure in the *top index* and IcH}t#}uJ}v4 is saved on line 420,904,771 in the main index for the first slice.

The information in two indices is stored in a B+ tree structure with main index as leaves and information in top index as internal nodes that contain offsets to all the blocks, the full 7-character prefix for the first 25-mer in the block, and the number of subjects for the 25-mer with the largest number of subjects in the block. The largest number is used during search to skip blocks for masking if the block has only one row, and the count is above the masking count threshold. The top index portion of the B+ tree is stored on a local solid-state drive for fast access while the leaves for the main index are on the network attached random access storage.

### Handling of metadata and suppressed subjects

To map encoded subject strings back to corresponding accessions, we maintain one translation table per slice that contains an offset for each subject in a *vocabulary* file. If metadata information is available for subjects, then that information is also incorporated into the vocabulary file. Some subjects that are present in the database may be suppressed later. To avoid rebuilding indexes with each such suppression, we zero-out the entries corresponding to the suppressed subjects in translation tables and the vocabulary file.

### Parameter determination

For each database, we compute default values for score constant and masking threshold parameters as follows. The score constant default is a round number near 0.4% of the number of subjects in the database where the rounding varies to the nearby tens or hundreds or thousands depending on the order of the value of the constant. The masking threshold default is the count such that ignoring all kmers with number of subjects more than this count removes 0.0005% of all harvested kmers in the database.

### Updating the database

The database design is modular – it supports multiple volumes in the database and splits each volume in 128 slices so that each file in the database is only a fraction of the total size of the database. Utilizing multiple volumes also makes it possible to have a large static volume that does not change with changes in the resource and a small dynamic volume that is updated at some frequency.

### Scoring

A score constant *C* is specified per database. For kmer *K*, let *K_N_* denote the number of subjects in the database from which *K* was harvested. The *kmer score* for *K* is calculated as *C/(C+K_N_)*. This calculation downweighs commonly occurring kmers; the higher the number of subjects a kmer is sampled from, the lower the score given to the kmer with the rate of downweighing determined by *C*.

For each query *Q*, the distinct kmers sampled from *Q* can fall in three mutually exclusive categories: the kmer is masked, the kmer is unmasked and was also sampled from at least one subject, and the kmer is unmasked and was not sampled from any subject. We will refer to these sets of kmers as K^masked^, K^matched^, and K^unmatched^, respectively.

The score computations do not consider kmers in K^masked^ – the size of this set is reported in “Show statistics” under the results on the web page. The following two values are computed per query: **Maximum raw score** *M* possible for *Q* is computed as the sum of kmer scores for all kmers *K* ∈ K^matched^ **Query match fraction** *F* for *Q* is ǁK^matched^ǁ / (ǁK^matched^ǁ+ǁK^unmatched^ǁ) Let K^S^ be the subset of K^matched^ that has all kmers with a match to subject S. For each query *Q* and each subject *S* with ǁK^S^ǁ > 0, following three values are computed:

**Raw kmer score** *R* for *S* on *Q* is the sum of kmer scores for all kmers *K* ∈ K^S^

**Percentage coverage** for S on Q is ǁK^S^ǁ *100 / (ǁK^matched^ǁ+ǁK^unmatched^ǁ)

**Pebblescout score** *P* for the pair *Q* and *S* normalizes raw kmer score for *S* on *Q* by the maximum raw score possible for *Q* and the query match fraction for *Q*. It is computed as *P = F * (R/M) * 100*

### Code and data availability

The metadata for all databases built, queries and output for all applications presented, the software for Pebblescout, an example for building a small database and searching the built database are available at https://ftp.ncbi.nlm.nih.gov/pub/agarwala/pebblescout/v2.25_release.zip. All runs and assemblies mentioned in the manuscript were publicly available at NCBI as of July 1, 2023. Access to the databases is through the search functionality available on the web page https://pebblescout.ncbi.nlm.nih.gov/

## Acknowledgements

We thank NCBI management for providing the resources needed to develop Pebblescout. Specifically, we thank Eugene Yaschenko for navigating the issues related to making the Pebblescout search publicly available and to Sergiy Ponomarov for developing the web page for the same. NCBI’s systems team, specifically Ron Patterson, was very helpful in providing access to tools for monitoring system loads. We thank David Lipman for his interest, suggestions for improving the manuscript, and introducing us to Andrew Fire and Caleb Lareau who tested and used the web page in their work, and to Sarah Preheim who suggested the microbial community application. We also thank Priyanka Ghosh, Barbara Robbertse, and Igor Tolstoy for taking an active interest as early users of Pebblescout, Alexander Souvorov for doing an independent assessment of alignment and assemblies for several runs, Valerie Schneider for providing extensive editorial comments on drafts of the manuscript, and Jacob Asherman for his comments on readability of the manuscript from a layman’s perspective.

## Author contribution

S.A.S. proposed the presented solution, designed, and implemented the software. R.A. managed the project, did testing, and found applications. Both authors contributed to building databases, data interpretation, and writing the manuscript.

## Competing interests

The authors declare no competing interests.

## Funding

This research work was supported by the National Center for Biotechnology Information of the National Library of Medicine (NLM), National Institutes of Health.

## SUPPLEMENTARY MATERIAL

### Datasets and Metadata

For each of the six databases, the metadata fields made available in the Pebblescout indices are shown in Table S1. The list of assemblies and metadata for RefSeq assemblies was extracted on April 22, 2022 from https://ftp.ncbi.nlm.nih.gov/genomes/refseq/assembly_summary_refseq.txt

The metadata for WGS sequencing projects was extracted from the information maintained for the corresponding assembly record in the Assembly database at NCBI; information for all assemblies in GenBank can be downloaded from https://ftp.ncbi.nlm.nih.gov/genomes/ASSEMBLY_REPORTS/assembly_summary_genbank.txt

The metadata for SRA runs was downloaded as XML using “esearch” and “efetch” utilities provided by NCBI. For example,

`esearch -db sra -query DRR092721 | efetch-format native > DRR092721.xml`

To generate MeSH terms for the Human RNAseq database, we parsed all text fields in the XML metadata available for the read sets at the level of EXPERIMENT_PACKAGE. Information in XML includes study title and abstract for the BioProject containing the run. MeSH preferred vocabulary terms (e.g., “T-Lymphocytes” instead of “T-Cells”) and subtrees for Anatomy [A], Bacteria [B03], Viruses [B04], Diseases[C], Chemicals and Drugs [D], Analytical, Diagnostic and Therapeutic Techniques, and Equipment [E], Phenomena and Processes [G], and Age Groups [M01.060] were used from the 2022 MeSH release. A small number of non-informative common MeSH terms, such as DNA, Algorithms, Standards, Research were ignored.

**Table S1:** Metadata in Pebblescout indexes. Phase 3 databases are from 1000 genomes project submissions.

| Database | Metadata fields |
| --- | --- |
| Metagenomic | BioProject, BioSample, Organism, CollectionDate, ReleaseDate, Title |
| WGS assemblies | BioSample, TaxId, ScientificName, Isolate |
| RefSeq assemblies | BioProject, BioSample, SpeciesTaxId, OrganismName |
| Phase3 Runs | BioSample, SampleName, Sex |
| Phase3 BioSamples | SampleName, Sex |
| Human RNAseq | BioProject, BioSample, ReleaseDate, Title, MeSH |

Next, we present some properties for the databases as reflected in the metadata.

The metagenomic database has 3,236,120 SRA runs. PRJEB11419, with sequences from the “American Gut Project”, provided the maximum number of runs from a single BioProject to the index (37,745 runs). Most of the assemblies in WGS sequencing project database are bacterial; the five taxa with the most assemblies are pathogenic to humans: *Salmonella enterica* (220,118 assemblies), *Escherichia coli* (142,912 assemblies), *Campylobacter jejuni* (48,915 assemblies), *Listeria monocytogenes* (45,208 assemblies), and *Salmonella enterica subsp. enterica serovar Enteritidis* (33,745 assemblies). Most of the assemblies in RefSeq database are also bacterial; the five taxa with the most assemblies are *Escherichia coli* (27,802 assemblies), *Staphylococcus aureus* (13,873 assemblies), *Klebsiella pneumoniae* (12,767 assemblies), *Salmonella enterica* (10,817 assemblies), and *Streptococcus pneumoniae* (8,744 assemblies). In the Human RNAseq database, the top five MeSH terms that appear in most BioProjects are Therapeutics (3,261 BioProjects), Tissues (3,062 BioProjects), Cell Line (2,689 BioProjects), “Sequence Determinations, RNA” (1,935 BioProjects), and Sex (1,392 BioProjects).

### Observations from the six databases

We present information from the databases we built to highlight various properties of the indexing method. The main resource requirement of a large network attached random access storage is satisfied at NCBI by its computing systems that have a VAST^20^ cluster.

### Even distribution of 25-mers in slices

The large WGS assembly JADMNL01 for Australian lungfish is 34.5 gigabases long. It contributes the maximum number of 25-mers to the WGS database at the count of 2.18 million. These are saved in the 128 slices of the index in a tight range within 0.22% from 16,978,996 to 17,016,017. Similarly, the largest number of 25-mers harvested from any SRA run in the Human RNAseq database comes from SRR16525168, where 8.67 million 25-mers were harvested; these are also saved in the 128 slices in a tight range within 0.16% from 67,707,010 to 67,813,163. Among all subjects from which at least one million 25-mers were harvested in any slice, we found that SRR15929127, a run in the Human RNAseq database, exhibited the largest percentage difference in range at 0.83% from 1,151,864 to 1,161,448.

### Harvesting memory footprint and verification

Harvesting took less than 4 gigabytes for almost all subjects, except for outliers with large contigs, such as, JADMNL01 mentioned in the previous paragraph. In millions of harvesting and verification runs for creating the six databases, we encountered a handful of discrepancies and corrected them by reprocessing the affected subjects. These bad executions were traced to two servers with unexplained intermittent and silent hardware faults.

### Encoding example and compression characteristics

We use the first slice (slice 000) of WGS index to illustrate the encoding and compression achieved by Huffman code. Aggregated 25-mer row `19nbIcH}t#}uJ}v4` has prefix `19nbIcH` for the 42-bits from the 25-mer and suffix `}t#}uJ}v4` as encoding for subjects DWTA01 (encoded by `}t#`), DWQO01 (encoded by `}uJ`), and DWUD01 (encoded by `}v4`). The length of the encoding in this slice is 2 for the 5,438 assemblies contributing the most 25-mers, 3 for 268,955 assemblies, 4 for 868,186 assemblies and 5 for the 1,622 assemblies contributing the least number 25-mers to the index. The total number of (25-mer, subject) pairs saved using encoding of length 2, 3, 4, and 5 is 6,162,324,912 (52.26%), 3,038,129,798 (25.76%), 2,592,604,352 (21.98%), and 37,816 (0.0003%), respectively. That is, over 52% of the kmer-subject pairs have subject identifiers encoded by two characters.

### Using byte-sized literals in the alphabet

During the merge operation, using byte-sized literals allows for each thread handling individual input to remap strings using the new mapping for subject identifiers and a very fast single thread to perform merging of kmers; the thread doing the merge can simply concatenate strings for subject identifier encodings for the same kmer. All indexes were created using 92 printable characters. We converted the metagenomic index to binary and saw savings of 10% in the index size. The 1000 Genome Phase 3 database by Runs saw a reduction of 25%, but we did not publish it as the primary purpose for including Phase 3 data was to show how the number of subjects impacted the index size.

### Suppression

Only the metagenomic database has suppressed subjects. These are runs that were publicly available at the time of indexing but were suppressed from the public repository before the release of the database. The number of such runs is 12,724 and is not included in the count of 3,236,120 indexed runs in the metagenomic database.

### Accessing SRA

For the 1000 Genomes Project, several attempts to download sequences for 265 runs (out of 37,106) were unsuccessful at the time of indexing, including the single run ERR3243126 for BioSample SAMN01761578. Our process attempts to download a run five times before flagging it as unavailable.

### False positive rate in results with a perfect Pebblescout score

Subsampling of queries by kmers can introduce situations where all sampled kmers match a subject, but there is no sequence or sets of sequences in the subject (contigs in an assembly or assembly of reads in a Run) that are or can be assembled into a sequence perfectly matching the query. To assess the false positive rate reported by Pebblescout (when it reports a perfect score but there is no perfect sequence match to the query,) and to assess how the false positive rate changes with query length, we collected three sets of one million distinct sequences of length 42, 100, 500, and 1000 bases. These sequences (queries) were randomly extracted from contigs in RefSeq assemblies. Next, we found all RefSeq assemblies (subjects) to which extracted queries have a full length 100% identity alignment. Comparing number of subjects reported by Pebblescout with a perfect score to the true counts, our empirical analysis shows that the false positive rate is 18.5% (3.3 billion reported with perfect score vs. 2.7 billion in assemblies), 10.9% (1.9 billion vs 1.7 billion), 5.8% (728 million vs 686 million) and 4.9% (497 million vs 473 million) for queries of lengths 42, 100, 500, and 1000, respectively.

The drop in percentage of false positives in the empirical results above is expected because the longer the query, the lesser the ambiguity about its presence in a subject due to more positions being sampled. We also note that, as per the guarantee of Pebblescout, there were no false negatives. That is, all subjects for each query expected to be found were reported as having perfect scores by Pebblescout.

### Output format for Profile search

Profile search prints one line for each 25-mer position sampled from the query. If the sampled 25-mer is also present in the database, the line printed contains the sequence for the 25-mer, starting position of the 25-mer in the query, and the number of subjects from which the same 25-mer was sampled. For sampled 25-mers with no matches, the line printed contains the sequence of the 42-mer from which the 25-mer was sampled and the starting position of the 42-mer in the query. If the same 25-mer is sampled from consecutive 42-mers, only the first 42-mer is printed. The counts reported can be an overestimate in case the database has suppressed runs as the list of subjects is not retrieved and cross-checked against the list of suppressed subjects.

Table S2 shows three rows from the profile of NC_045512.2 using WGS and metagenomic databases. The first row using the WGS database is an example of a sampled 25-mer where no assembly has the 42-mer printed. The remaining rows show the variability in counts at different positions.

**Table S2:**
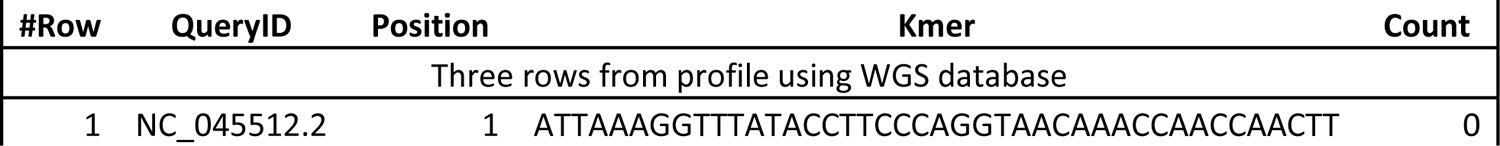
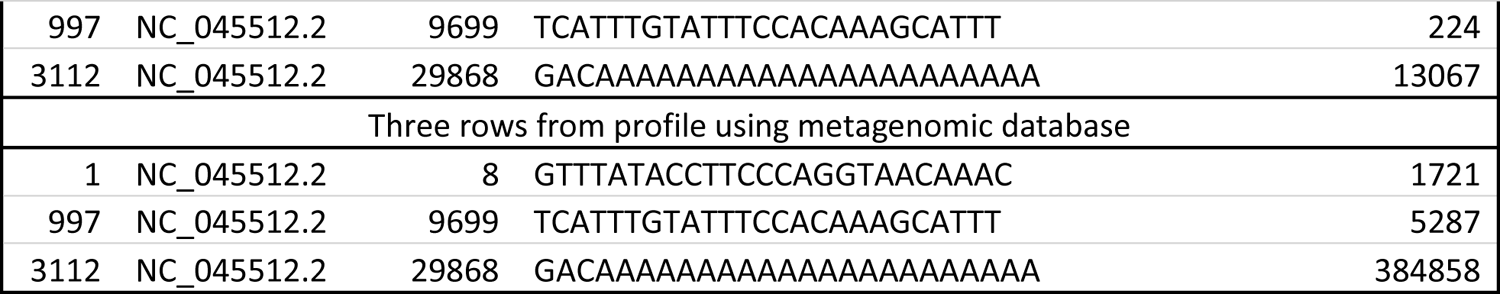
Sample for profile search output format

### Output format for Summary search

For each query and subject with a match for at least one kmer sampled from the query that is not masked, one row is present in the output of summary search. In addition to the query and subject identifiers, and three computed scores (raw score, percent coverage, and Pebblescout score), any additional metadata fields printed by default depend on the database. Pebblescout supports a user specifying which of the available metadata fields are printed, although this feature is currently not supported on the pilot web page. For each query, the rows are sorted by the Pebblescout score in descending order from the largest score to the smallest, and all such rows are consecutive in the output. Table S3 shows some rows from the Summary search using NC_045512.2 as query to the WGS database.

**Table S3:**
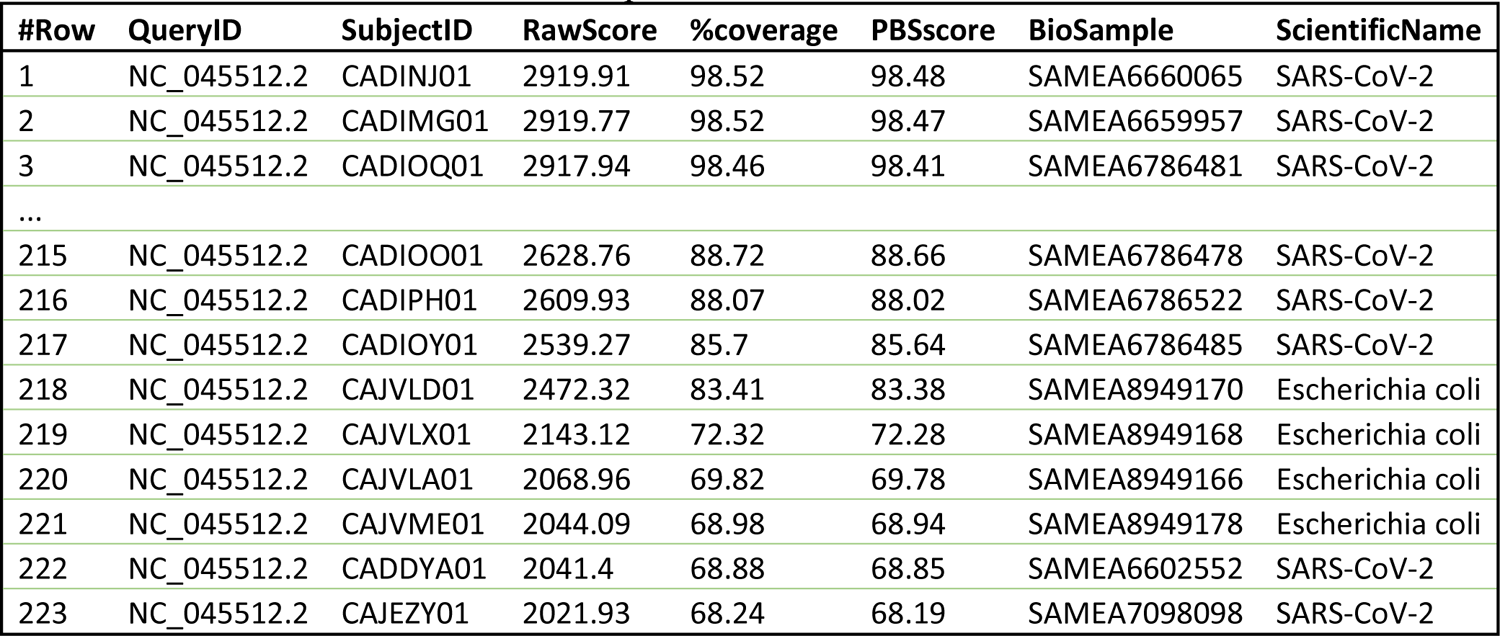
Sample for summary search output format. We abbreviated “Severe acute respiratory syndrome coronavirus 2” to “SARS-CoV-2” to save space in the column for ScientificName.

### Output format for Detailed search

Detailed search output has columns for query and subject identifiers, three scores, and metadata as in the Summary search output. In addition, it has columns for position, kmer, and count. Subject identifier has a dash when the 25-mer sampled has no matches in the database; as in profile search output, the first 42-mer from consecutive 42-mers sampling the same 25-mer position is printed. Subject identifier has an asterisk if the 25-mer sampled was masked. For all query-subject pairs reported, rows are grouped for each pair and sorted numerically by the position of sampled kmers in the query. Rows with a dash or asterisk are at the end of the output. These rows are not sorted by the printed position but by the position of the 25-mer; the starting position of the 42-mer may be less than the starting position of the 25-mer in the query.

Table S4 shows several positions less than 25 bases apart with a match between *Cyclospora cayetanensis* transcript XM_026338611.1 and SRR7686101 in the metagenomic index. The table does not show the query identifier and three columns for the three scores. Several positions with a count of 1 also show that the corresponding 25-mer was sampled *only* from SRR7686101 in the metagenomic database.

**Table S4:**
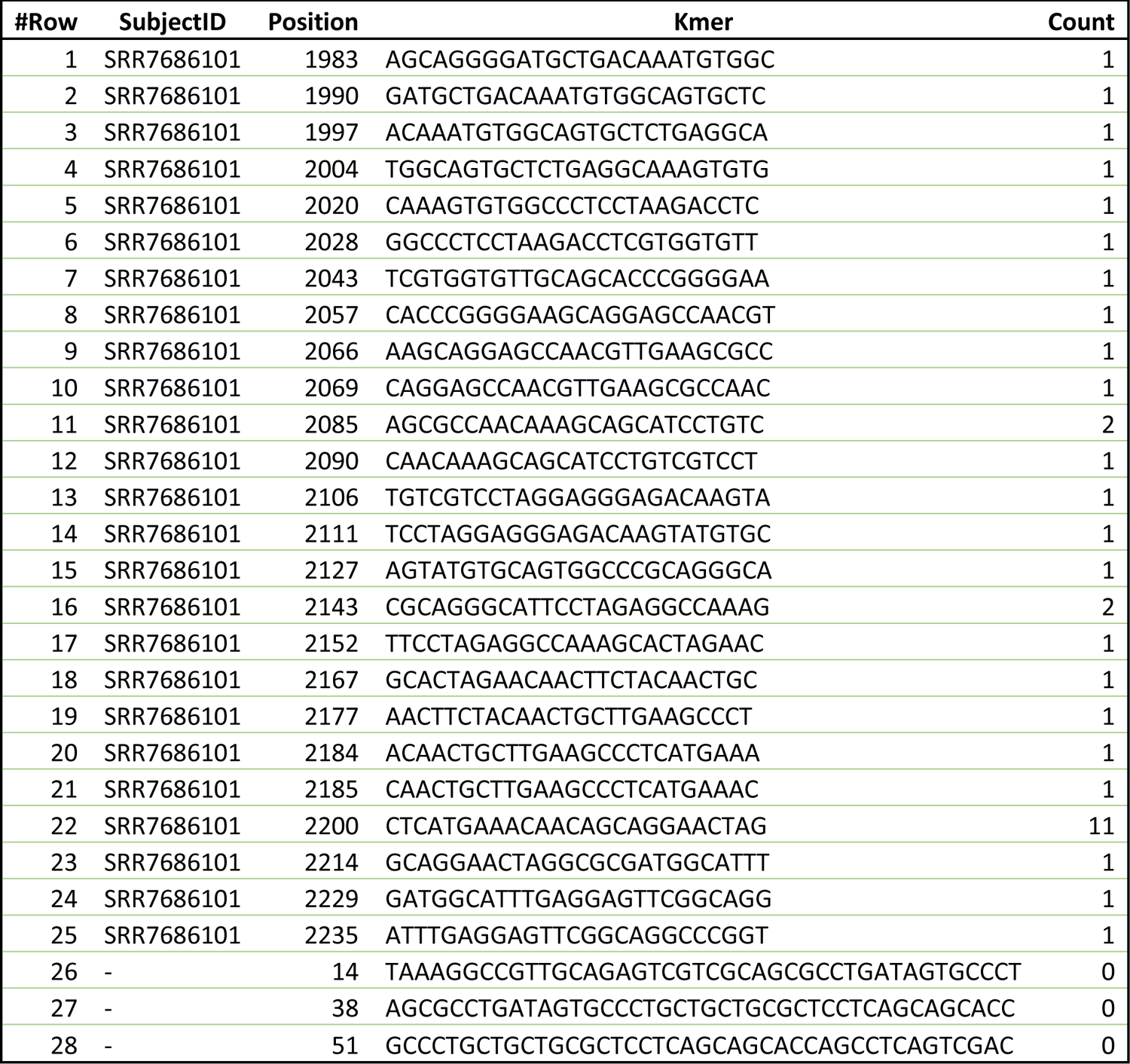
Sample output from Detailed search with columns for query identifier and three scores removed

### Analysis for search applications

We present detailed analysis for all ten applications below. Some of the information regarding queries and aim of the studies is also present in the main manuscript but is repeated here for the sake of clarity and completeness.

### Find previously undiscovered species related to known species (viral discovery example)

To our knowledge, the largest petabyte-scale sequence alignment to SRA was done for viral discovery using the Serratus pipeline^5^. In this analysis, the nucleotide search component of the pipeline aligned 12,295 query sequences Q (referred to as ‘cov3m’ reference set in [5]) to 3,837,755 runs R^Ser^ in ca. May 2020; the list of runs was generated using metadata from metagenomic, genomic, and RNAseq runs (Supplementary table 1(a) of [5]). Of these, 14,304 runs satisfied alignment filtering criteria used by the authors and were considered as good runs for use as inputs to genome assembly, referred to as R^Ngood^ below. Authors also identified additional runs for use in assembly by protein search, and by using species information available in SRA using STAT^7^ in January 2021. The total number of runs assembled using coronaSPAdes^18^ was 56,511 (file aindex.tsv in data for [5]). Assemblies for 11,120 runs were deemed good for further analysis by the authors, referred to as R^final^ below.

The overlap between R^Ser^ and runs present in Pebblescout metagenomic database R^Pbs^ is shown in Figure S1. The figure also shows distribution for R^Ngood^ in the blue oval, runs reported by Pebblescout summary search using Q against the metagenomic database in the grey rectangle with rounded edges, and distribution for R^final^ in the brown rectangle. We make the following observations:

1. Pebblescout search filters out 1,395,194 runs from the 1,653,559 runs (84.4%) present in both R^Seq^ and R^Pbs^ and 1,368,892 from the remaining 1,582,561 runs (86.5%) present only in R^Pbs^.
2. All 1,815 runs present in R^Ngood^ and R^Pbs^ are reported by Pebblescout. That is, using Pebblescout search to filter metagenomic runs would not have caused any loss of viral discovery reported in the study.
3. All but one run present in R^final^ and R^Pbs^ are reported by Pebblescout. The only run not reported by Pebblescout is SRR7408078. For this run, the authors reported only one contig assembly of length 461 bp. The full length of the contig has 95% identity match to CP027235.1 (*Haemophilus sp.*), a bacterial species, so the reported sequence is unlikely to be a viral sequence.

Next, we looked for viral sequences in runs found by Pebblescout that are neither in R^Ngood^ nor R^final^. We restricted our search for examples to runs submitted to SRA before May 2020 from runs present in R^Ser^ (set of 256,550 in the figure) and before January 2021 for runs not in R^Ser^ (set of 213,001 in the figure) to match the dates reported in the study^5^. Examples we found for the two categories include SRR6434387 from BioProject PRJNA428222, and all 43 runs from BioProject PRJNA633210, respectively.

We assembled SRR6434387 using coronaSPAdes. The assembly has contigs that appear to be novel coronavirus sequences. For example, four contigs in SRR6434387 assembly align to 80.6% of the feline spike protein (AB535528.1 from 75 to 3743 bp) with alignment percent identities ranging from 77% to 91.8%, demonstrating that SRR6434387 has viral sequences. Furthermore, five runs other than SRR6434387 from PRJNA428222 are in R^final^. It is interesting to note that SRR6434387 sequenced sample for intestinal viral and bacterial microbiota of mink from a farm in Denmark and SARS-CoV-2-positive cats and dogs were previously suspected of transmitting the virus to minks in several studies, including [21] that states mink farms in Netherlands were culled to prevent transmission.

The study^22^ that submitted PRJNA633210 also submitted virus assemblies from these runs in GenBank under accession numbers OL770284 to OL770294. These assemblies include Human coronavirus sequences.

Hence, we conclude that Pebblescout reported all good runs identified by the authors of the study^5^, potentially identified several additional runs of value, and did so while substantially reducing the number of runs that were aligned in one step of the Serratus pipeline.

**Figure S1:**
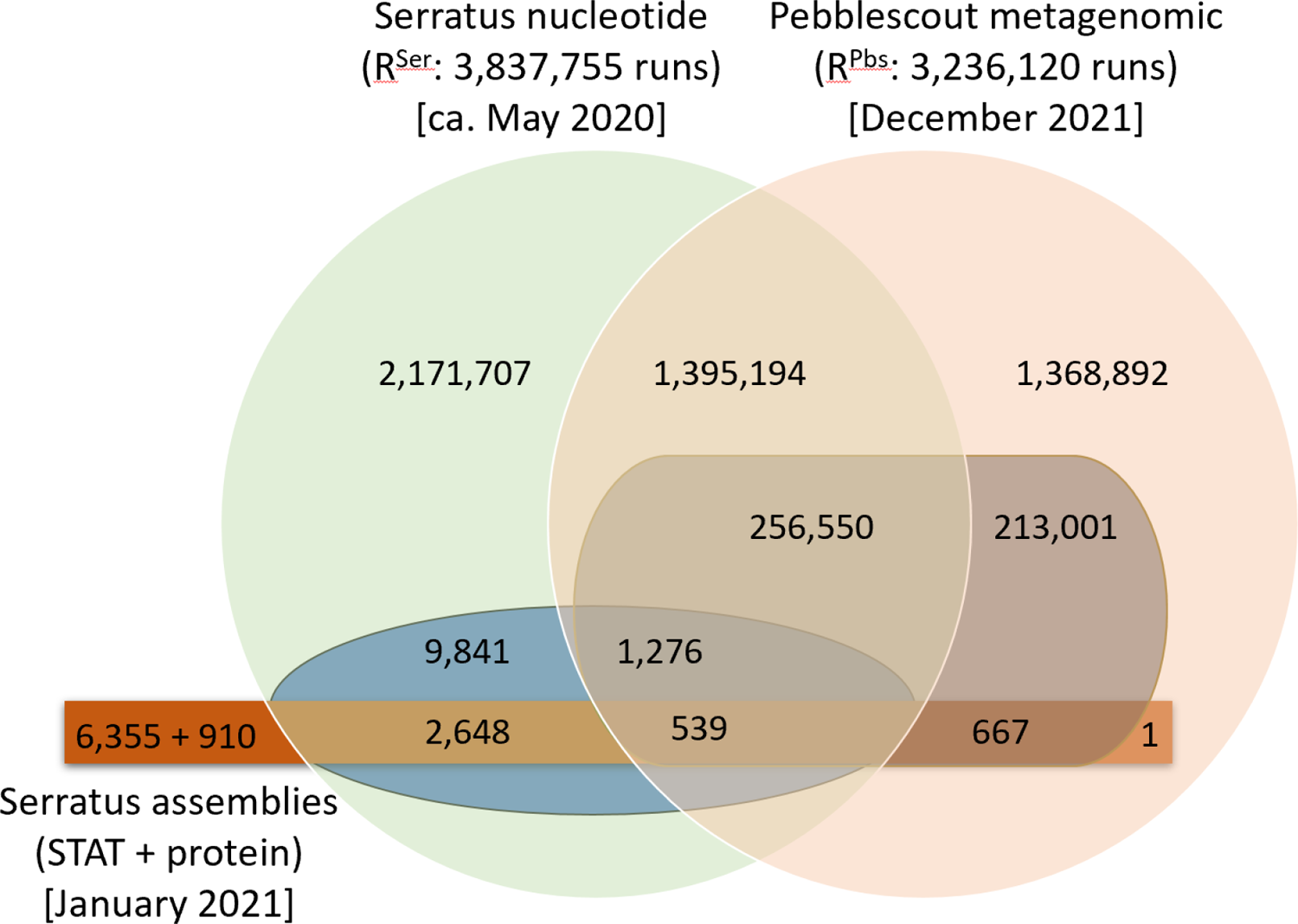
Total count of runs aligned in the nucleotide search component of the Serratus pipeline used for viral discovery (R^Ser^; green circle) and runs present in the Pebblescout metagenomic index (R^Pbs^; peach circle). The blue oval shows distribution of 14,304 runs from the nucleotide search that were considered as good for assembly by the Serratus pipeline (R^Ngood^) while the grey rectangle with rounded edges shows runs reported by Pebblescout. The brown rectangle shows distribution of 11,120 assembled runs considered by the authors of the study^5^ for further analysis (R^final^). This set includes 910 runs reported by protein search and 6,335 found by STAT; for runs reported by more than one method, we assigned the runs to nucleotide set first, then protein, and rest to STAT.

### Find known species in new environments (*Candida auris* example)

A recent study^6^ took a metabarcoding approach to identify an environmental niche of *Candida auris*. The authors aligned the ITS region (ITS1+5.8S+ITS2) of *C. auris* as query (accession NR_154998.1 from 1 to 320 bp) to 603,692 runs. The metabarcoding and metagenomic runs containing fungal ITS in SRA were identified using keywords “fungi, ITS1, ITS2, mycobiome” and whole genome runs were excluded from the list. The list was created between January and March 2021, and the alignment was done using BLAST with a percent identity threshold of 99%. The analysis found *C. auris* in 34 runs. Only 19 of these 34 runs are in our Pebblescout metagenomic database as the remaining 15 runs are tagged as “genomic” in the SRA metadata.

To find a match count threshold that separates species-specific kmers from other species, we did a profile search to the metagenomic index using the same query used by the authors of the study^6^. The profile reported 29 sampled kmers with match counts distributed as follows: 12 kmers with the match count of at most 76, followed by 11 kmers with counts in the range from 357 to 20,810, and remaining 6 kmers with match counts at least 143,583. All kmers sampled from the species-specific ITS2 region of the query have match count at most 76 while all kmers with match count 143,583 or higher are from the non-species-specific 5.8S region. These observations and jumps from 76 to 357 and from 20,810 to 143,583 suggest (i) using a masking threshold of 76 to retain kmers that are specific to *C. auris* and (ii) supporting the choice of 76 by analyzing the distribution of species using sampled kmers with a match count of at most 20,810.

To find the distribution of species, we aligned twenty-three kmers with match count of at most 20,810 sampled by Pebblescout from the *Candida auris* ITS sequence to the nucleotide non-redundant database using BLAST with word size 25. Next, we used the description for the matching sequences in GenBank to assign a species to each subject. Table S5 shows the kmer, match count in metagenomic index, and the distribution of species found. We note that the same sequence from *Candida haemuloni* matches all kmers with profile count at most 357. This sequence is KX420743.1 that is likely misidentified and is a *Candida auris* sequence.

**Table S5:**
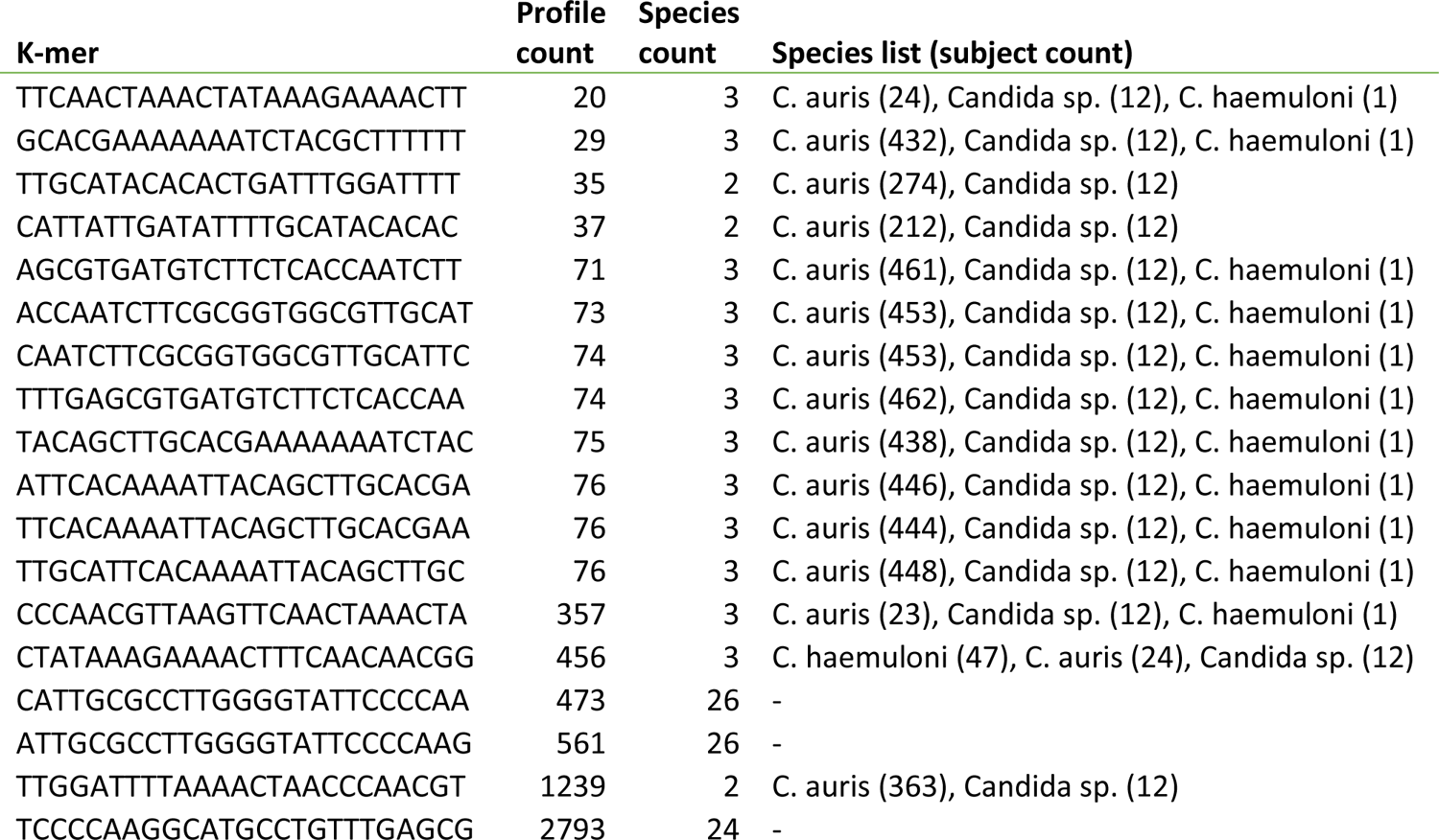
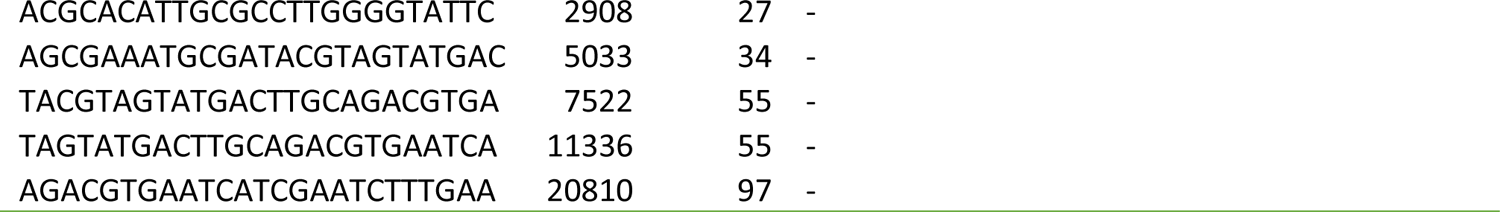
Kmers sampled by Pebblescout from the *Candida auris* query, match counts in profile using metagenomic database, number of species matching the kmer in nucleotide non-redundant database, and list of matching species. For each species, number of sequences from the non-redundant database that have a match is listed in the brackets. We did not list species when number of species for a kmer is more than 20 <u>and use a dash instead.</u>

Using a masking threshold of 76, a Pebblescout summary search of the metagenomic database using the same query that authors used in the study [6] finds 129 runs, including all 19 runs identified in the published study present in the metagenomic database. Aligning the ITS1 and ITS2 regions of the query sequence to these 129 runs using BLAST and looking for reads that align to the full length of ITS1 or ITS2 at >=95% identity shows that such reads are present in 86 runs. However, even at this reduced percent identity stringency compared to the published work, 7 of the 19 previously identified runs are not included in these 86 runs -- we must allow nine bases at one end of the query to remain unaligned to find an alignment for these 7 runs. We note that the computational effort needed to analyze 129 runs by alignment is significantly smaller compared to doing the same with 603,692 runs.

From the 86 runs identified above, excluding all runs submitted in 2021 as some of them may not have been available at the time of the original study, excluding runs reported in the original study, and excluding runs that have metadata in their associated BioProject records indicating they come from studies that are associated in some way with *C. auris* specifically or other fungal associated samples, we have 7 runs from three BioProjects that are likely of high interest as environmental niches for *C. auris*: two from PRJNA488992 (SRR8584355, SRR8584356: wastewater drains and river), one from PRJNA610489 (SRR11241457: soil sediment from India), and four from PRJNA666746 (SRR12750709, SRR12750710, SRR12750711, SRR12750712: Dysbiosis of Oral Microbiota during Oral Squamous Cell Carcinoma Development). Hence, we conclude that use of Pebblescout for this study, had it been available at the time, would not only have identified the same runs of interest, but would have reduced the computational effort required and would have returned additional runs that would likely have been of interest.

### Find host species for endosymbionts (*Wolbachia* example)

A recent study^12^ used 44 reference transcripts for three genes (*ftsZ, groE, wsp*) from 36 host species to identify additional arthropod host species for *Wolbachia*. The authors found 2,545 runs from 288 host species in SRA using various metadata criteria available in “Run Selector” provided on the SRA web page. They downloaded at most 5×10^7^ reads for each run and aligned them to the reference transcripts using Magic-BLAST^23^. They also used an iterative bait and assemble strategy for assembling genomes using seven reference genomes representing *Wolbachia* isolates from insects and nematodes.

We used Pebblescout summary search using the same 44 reference transcripts as queries to the metagenomic database. Because we wished to recover full-length genes, we only considered subjects with a Pebblescout score of >=90 for at least one query. We further limited the subjects to only those runs that: (i) have a host species specified; (ii) the host is an anthropod species; (iii) the species is not one of the 36 host species for any reference transcripts used as queries; and (iv) the species is not in the list of 288 host species considered by the authors in their analysis. We found 35 runs from 16 species that passed all criteria. For each of the 35 runs, we subsequently performed genome assembly using SKESA^24^ and reference-based gene assembly using SAUTE^25^. We used BLAST to align these assemblies to the reference transcripts and to all seven reference genomes used by the authors in their assembly pipeline. This analysis found that for all 16 species we identified, there is at least one run whose assembly has a contig to which at least one reference transcript has a full-length alignment. Coverage in alignments using contigs in genome assemblies for the 16 species to reference genomes ranges from 14 Kb to 1.34 Mb; this translates to 1% to 90% of the corresponding reference genome as the length of NC_010981.1 and CP001391.1 is 1.48 Mb and 1.44 Mb, respectively. For each host species identified, Table S6 shows the run which produced the maximum coverage to a reference genome from all runs for the same species. It also shows genes for the same run that have alignment to the full length of a reference transcript.

**Table S6:**
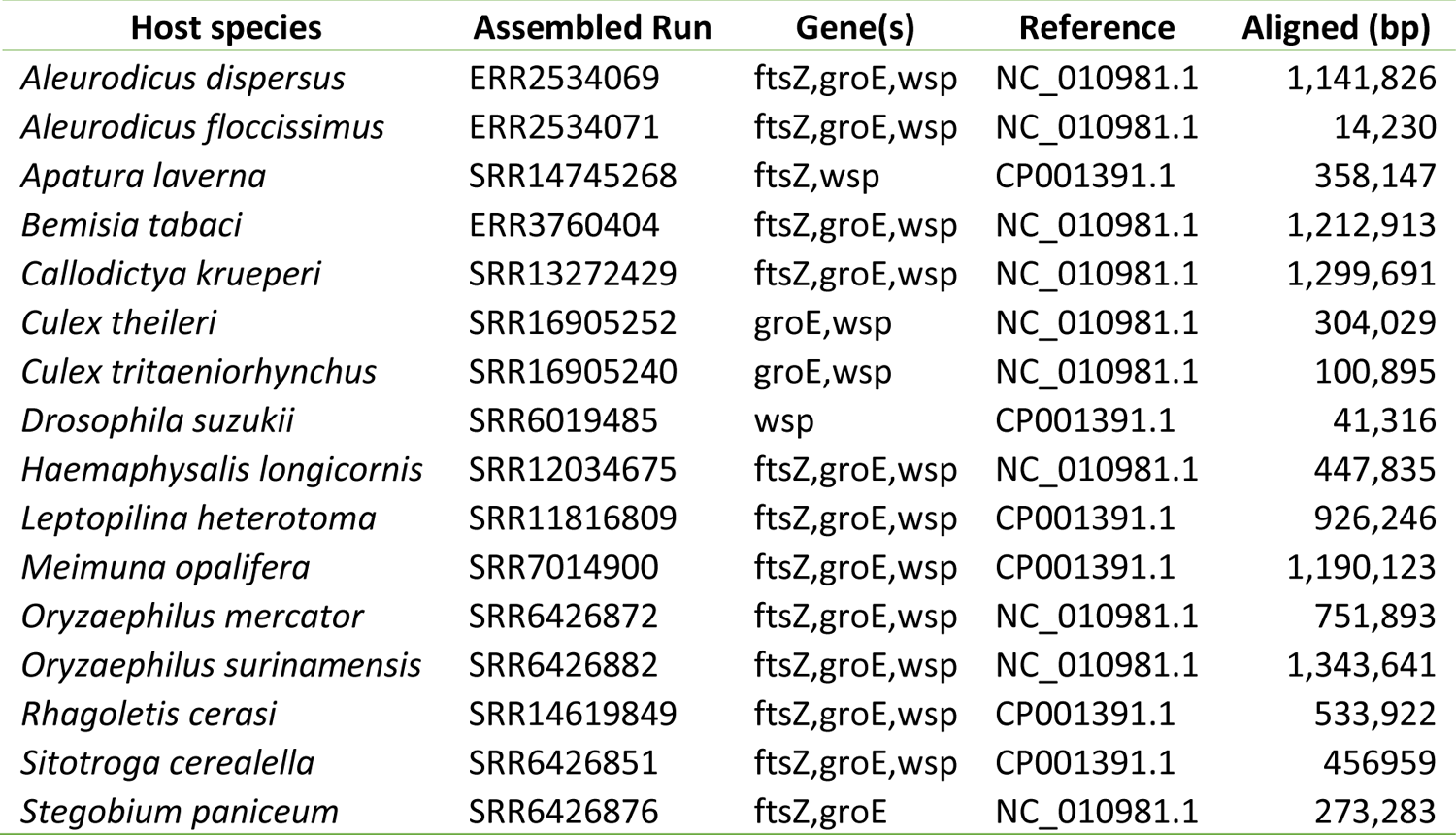
*Wolbachia* host species and SRA run leading to the discovery of the host species

| Host species | Assembled Run | Gene(s) | Reference | Aligned (bp) |
| --- | --- | --- | --- | --- |
| *Aleurodicus dispersus* | ERR2534069 | ftsZ,groE,wsp | NC_010981.1 | 1,141,826 |
| *Aleurodicus floccissimus* | ERR2534071 | ftsZ,groE,wsp | NC_010981.1 | 14,230 |
| *Apatura laverna* | SRR14745268 | ftsZ,wsp | CP001391.1 | 358,147 |
| *Bemisia tabaci* | ERR3760404 | ftsZ,groE,wsp | NC_010981.1 | 1,212,913 |
| *Callodictya krueperi* | SRR13272429 | ftsZ,groE,wsp | NC_010981.1 | 1,299,691 |
| *Culex theileri* | SRR16905252 | groE,wsp | NC_010981.1 | 304,029 |
| *Culex tritaeniorhynchus* | SRR16905240 | groE,wsp | NC_010981.1 | 100,895 |
| *Drosophila suzukii* | SRR6019485 | wsp | CP001391.1 | 41,316 |
| *Haemaphysalis longicornis* | SRR12034675 | ftsZ,groE,wsp | NC_010981.1 | 447,835 |
| *Leptopilina heterotoma* | SRR11816809 | ftsZ,groE,wsp | CP001391.1 | 926,246 |
| *Meimuna opalifera* | SRR7014900 | ftsZ,groE,wsp | CP001391.1 | 1,190,123 |
| *Oryzaephilus mercator* | SRR6426872 | ftsZ,groE,wsp | NC_010981.1 | 751,893 |
| *Oryzaephilus surinamensis* | SRR6426882 | ftsZ,groE,wsp | NC_010981.1 | 1,343,641 |
| *Rhagoletis cerasi* | SRR14619849 | ftsZ,groE,wsp | CP001391.1 | 533,922 |
| *Sitotroga cerealella* | SRR6426851 | ftsZ,groE,wsp | CP001391.1 | 456,959 |
| *Stegobium paniceum* | SRR6426876 | ftsZ,groE | NC_010981.1 | 273,283 |

This analysis demonstrates that Pebblescout can be used to create an initial list of subjects using the sequence information that can be subsequently filtered using the study criteria. Doing so the other way around, as was done in the published study, leads to missing some interesting subjects and an incomplete analysis of the data publicly available in SRA.

### Find assemblies in WGS for additional context or to support novel sequence findings (ssRNA phage example)

A study^19^ published in Feb 2020 to expand the set of known ssRNA phage genomes identified 15,611 novel sequences. It appears to us that authors compared their sequences to previously known ssRNA phage sequences to claim that the sequences they assembled are novel but, understandably, did not check the novelty with respect to sequences in WGS due to lack of publicly available easy to use tools to do so.

Using these 15,611 sequences as queries, Pebblescout finds one WGS assembly, UYOA01, with Pebblescout scores of >=25 for at least a thousand queries. Assembly UYOA01 was released a year before Feb 2020. BLAST alignments of the query sequences to UYOA01 finds 388 queries that have a full-length alignment with ≥95% identity to contigs in UYOA01. This example shows that Pebblescout provides an easy and effective way to find assemblies in WGS that have contigs with similar sequence as the given query. Such an analysis can provide additional context for the new sequences or show that nothing similar exists in WGS.

### Detect hard-to-sequence samples (*Cyclospora cayetanensis* example)

*Cyclospora cayetanensis* is an apicomplexan pathogen that is highly host-specific to humans and causes foodborne disease outbreaks^26^. It can be detected within feces of infected individuals but identifying it in the environment is highly challenging^27^. It is also not known if there are other species closely related to *C. cayetanensis.* Therefore, it is important to find *any* reservoirs for the genome with data in SRA so we can learn more about *C. cayetanensis* and related species. As finding such sequences is rare, subjects with matches to only small regions of queries are more likely than those with full length matches. Distinguishing spurious short matches from those that are meaningful requires additional filtering, such as by identifying maximal regions of the query where every base is in a sampled kmer with a match to the subject; such regions of length >=100 bases are hereafter referred as *partial matches* to the subject. Position information for matching kmers per subject is present in the output of the Pebblescout detailed search, therefore facilitating the identification of such partial matches.

Using 5,793 RefSeq mRNA sequences annotated on the *C. cayetanensis* assembly (accession GCF_002999335.1) as queries, we performed a Pebblescout detailed search of the metagenomic database. The output has 14,254,556 query-subject pairs from 365,582 subjects with any match. Filtering for partial matches reduces the output to 6,541 query-subject pairs from 1,540 queries and 2,831 subjects. We aligned all 6,541 pairs using BLAST with options for word size of 25 and no low complexity filtering, identifying 6,330 pairs with a >=100 bp alignment between the transcript and a read in the subject.

Further restricting the BLAST results to query-subject pairs where at least five reads have a >=100 bp alignment at >=97% identity to the transcript reduces the number of pairs to 165. We observe that 70% (n=116) of these pairs involve reads from SRR12874575. In the remaining 49 pairs, a different SRA run is present in each pair, but the query transcript XM_026335942.1 is implicated in 42 (85%) of them. The remaining seven pairs are shown in Table S7. Additional analysis of the BLAST alignments for these 7 pairs shows that the two runs with over 250 reads each aligned do so to a very small region of the transcript while two runs SRR11414099 and SRR14160768 with only a small number of reads aligned do so at much higher coverage (>=50%). This suggests that total count of aligned reads alone may not be the best measure for the determining the presence of a query in the run.

An example of a run in the set of 6,330 pairs but not in the set of 165 pairs is SRR7686101. It is in three pairs with partial matches but number of reads aligning for the three pairs is four for one and one each for the other two. However, all six alignments have percent identity >=99.1%. While further analyses are required, alignments for SRR7686101 suggest that other runs in the originally identified set of 6,330 query-subject pairs may come from samples containing *C. cayetanensis*. Thus, Pebblescout may prove to be a particularly valuable tool for the identification of reservoirs containing incomplete genomes, such as those associated with difficult-to-sample species.

**Table S7:**
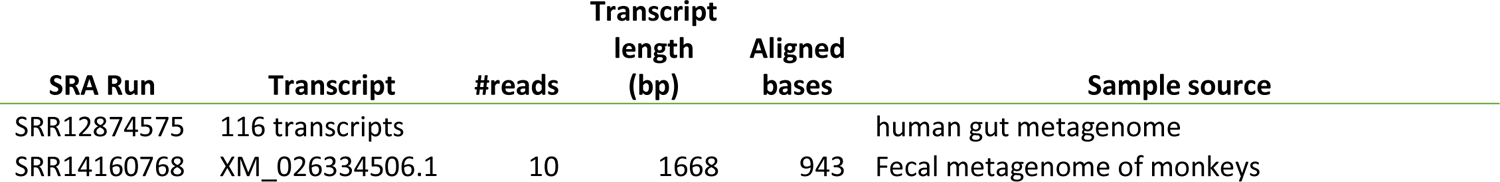
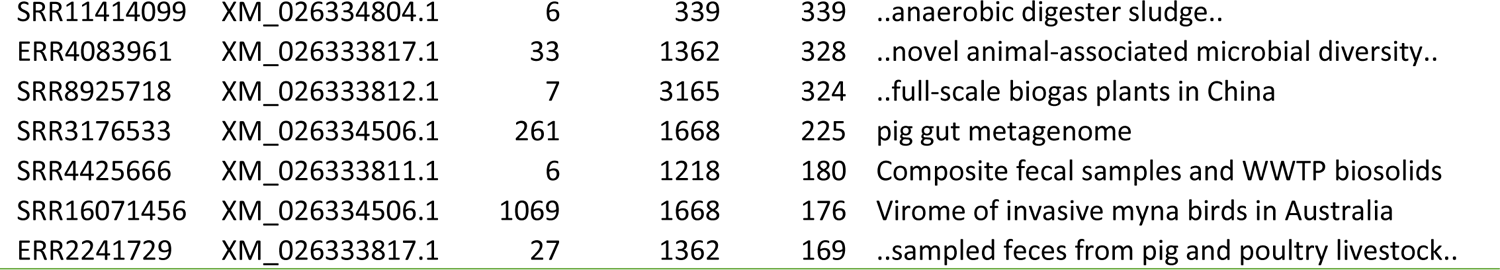
SRA runs and their sources with partial matches to *Cyclospora cayetanensis* transcripts. Count for reads that align to the transcript for >= 100 bases at >=97% identity is in (#reads)

### Find variant/invariant regions of genomes (*E. coli* and HIV-1 examples)

Assuming all kmers are equally likely to be sequenced, the kmers with low counts in the profile search output represent positions of high variability compared to kmers with high counts. We present our findings from the profile search using genomes for the most well studied strain in laboratory *E. coli* K12, and one of two known viruses that cause AIDS, HIV-1, queried against metagenomic and Human RNAseq databases, respectively.

An *E. coli* K12 (accession NC_000913.3) profile search of the metagenomic database finds an average of 54,137 matches and median of 54,837 matches for the 489,474 kmers sampled from the query. The results have five sampled positions with no more than 250 matches. These query positions are 3,424,218, 3,424,227, 4,296,368, 4,296,373, and 4,296,377 with match counts of 50, 55, 70, 28, and 12, respectively. Hence, these five positions, seen rarely in the metagenomic database, may be positions that evolve rapidly or have some other significance. Knowledge of such positions may be useful in designing signatures for pathogen detection^28^ or for understanding evolution of strains over time^29^.

Human RNAseq database is not expected to have many subjects infected by HIV-1. However, as RNAseq is an unbiased way of detecting viruses^30^, we assessed the profile for the HIV-1 genome (accession NC_001802.1) queried against the Human RNAseq database, shown in Figure S2, to highlight variability seen in the database. Position 4,118 matched only one subject. Six positions with no matches are bases 4,119, 4,120, 4,121, 4,125, 5,598, and 6,671. Figure S2 shows that remaining kmers are well separated using match count of 512 (red bar in Figure S2) and 4,096 (blue bar in Figure S2).

Using kmers reported in the profile output with counts higher than 4,096, doing BLAST for these kmers to non-redundant nucleotide database with word size 25, removing matches to HIV-1 and synthetic constructs, and counting subjects that have matches to almost all high-frequency kmers shows that all such matches are due to cloning vectors, with Lentiviral as the most frequent description. Hence, we posit that the high-frequency positions above the blue bar are likely due to cloning vectors, such as Lentiviral vector SCRPSY (accession KT368137.1), used in replication.

Positions below the red bar are likely positions with higher variability in HIV-1 compared to the positions between the red and blue bars. Support for this hypothesis comes from low counts in the sequence range outlined by the black box for the gp160 region that is known to be variable^31^.

**Figure S2:**
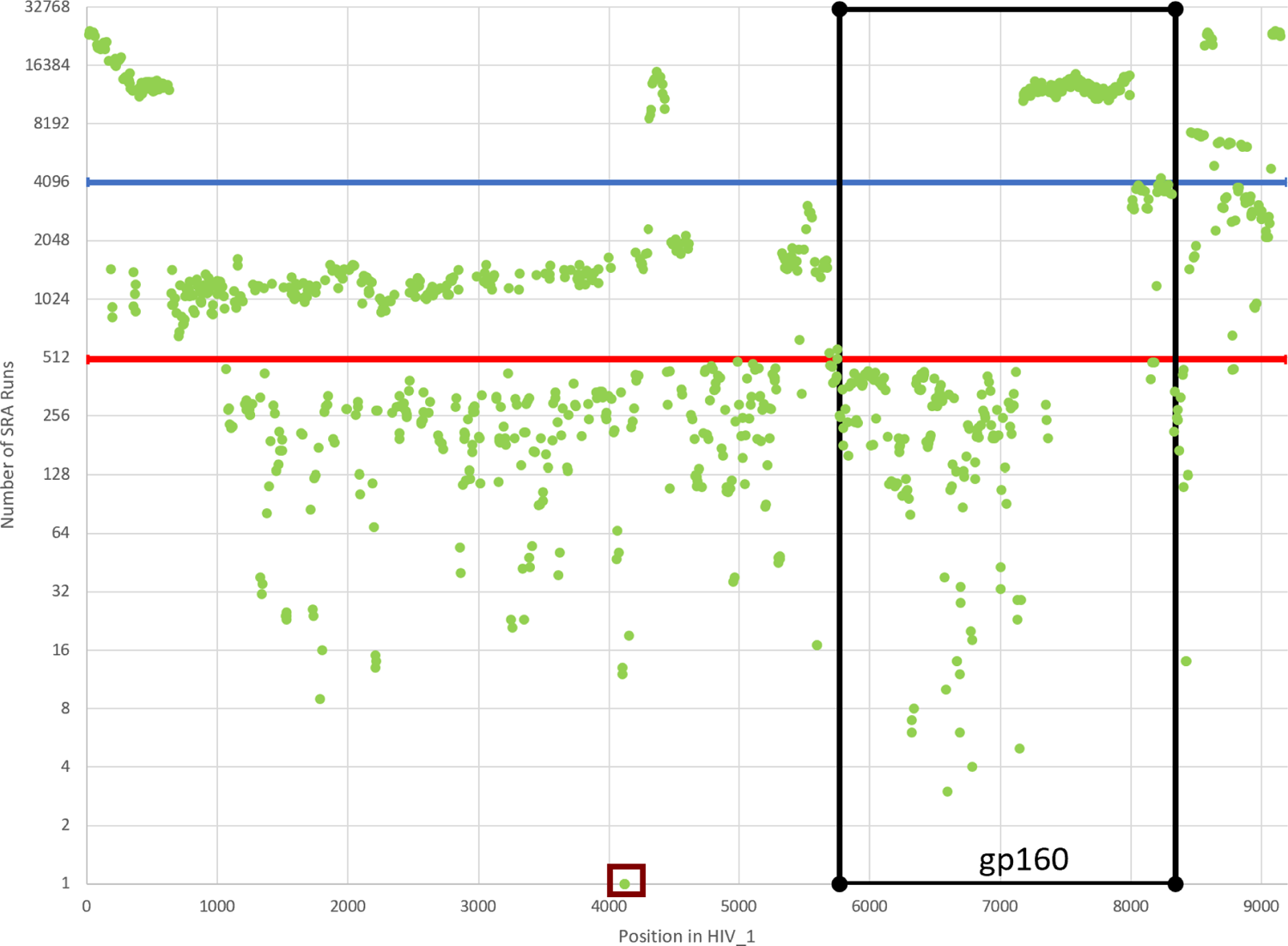
HIV-1 profile against the Human RNAseq index. Position on the HIV-1 genome is shown on the X-axis. Y-axis shows the number of subjects with a match in the Human RNAseq database for kmers sampled at various positions.

### Find contamination (SARS-CoV-2 example)

Using SARS-CoV-2 genome (accession NC_045512.2) as the query and searching the WGS database, we find that the top 217 matches are to SARS-CoV-2 genomes, followed by four *E. coli* genomes. Part of this output is shown in Table S4. All WGS genomes with a Pebblescout score of at least one to WGS are either SARS-CoV-2 assemblies or *E. coli* assemblies. There are 36 such *E. coli* assemblies; all are from BioProject PRJEB46014. Aligning these *E. coli* genomes to SARS-CoV-2 genome and comparing coverage to Pebblescout score gives R^2^ of 0.99862; Pebblescout scores are in the range from 1.76 to 83.38.

### Find genes of interest (Methicillin-resistance genes example)

Methicillin-Resistant *Staphylococcus aureus* is a superbug of worldwide concern^32, 33^. mecA is the most common gene mediating methicillin-resistance, but homologues mecB, mecC, and mecD were discovered recently: mecB in *Macrococcus caseolyticus* that colonizes animal skin, mecC in livestock which has since spread into clinical isolates, and mecD in bovine and canine *Macrococcus caseolyticus* isolates. Table S8 shows the number of subjects in each database that have a perfect Pebblescout score for the corresponding gene variant. The search used all kmers by removing masking limit; default masking limit removes up to 94% of kmers for some mecA variants.

**Table S8:** Number of subjects with perfect Pebblescout score to MRSA transcripts in RefSeq, WGS, and metagenomic databases

| Gene | Accession | Length (bp) | RefSeq | WGS | Metagenomic |
| --- | --- | --- | --- | --- | --- |
| mecA.1 | NG_047937.1 | 2207 | 0 | 0 | 13 |
| mecA.2 | NG_047939.1 | 2207 | 1010 | 1919 | 1162 |
| mecA.3 | NG_047941.1 | 2007 | 1 | 0 | 6 |
| mecA.4 | NG_047940.1 | 2007 | 0 | 0 | 86 |
| mecA.5 | NG_047938.1 | 2207 | 2 | 0 | 111 |
| mecA.6 | NG_047945.1 | 2107 | 24 | 37 | 31 |
| mecA.7 | NG_047946.1 | 2007 | 0 | 0 | 0 |
| mecA.8 | NG_047947.1 | 2007 | 0 | 0 | 6 |
| mecA.9 | NG_047948.1 | 2007 | 0 | 0 | 0 |
| mecA.10 | NG_047936.1 | 2207 | 0 | 0 | 0 |
| mecB.1 | NG_047953.1 | 2225 | 2 | 1 | 7 |
| mecB.2 | NG_047954.1 | 2225 | 1 | 0 | 0 |
| mecC | NG_047955.1 | 2198 | 0 | 0 | 0 |
| mecD | NG_054960.1 | 2237 | 36 | 26 | 0 |
| Total |  |  | 1076 | 1983 | 1422 |

Upon aligning each of the 1,076 query-assembly pairs with a perfect Pebblescout score to the RefSeq database using BLAST, we find that all of them have 100% identity alignments for the entire length of the corresponding query – 1070 pairs have a single full-length alignment, 3 pairs have two full length alignments (mecA.2 against GCF_000641435.1, GCF_002089055.1, GCF_003239745.1), and 3 pairs have alignments split in contigs. Pairs with split alignments are: (i) mecA.2 alignments to GCF_000683395.1 split on NZ_JILJ01000024.1 and NZ_JILJ01000030.1, (ii) mecA.2 alignments to GCF_014637435.1 split on NZ_JACJBQ010000097.1, NZ_JACJBQ010000417.1, and NZ_JACJBQ010000064.1, and (iii) mecD alignments to GCF_019357515.1 split on NZ_CP079950.1 and NZ_CP079950.1

By aligning each of the 1,983 query-assembly pairs with a perfect Pebblescout score to the WGS database, we find that all of them have 100% identity alignments for the entire length of the corresponding query – 1541 pairs have a single full-length alignment, 438 pairs have two full length alignments, mecA.6 against DADKIB01 has three full length alignments, and 3 pairs have alignments split in contigs. Query in all three split alignment pairs is mecA.2: alignments to (i) JACJBQ01 are split on three contigs: JACJBQ01.64, JACJBQ01.97, and JACJBQ01.417, (ii) JAEVMO01 are split on four contigs: JAEVMO01.66, JAEVMO01.106, JAEVMO01.215, and JAEVMO01.573, and (iii) JILJ01 are split on two contigs: JILJ01.24 and JILJ01.30.

For metagenomic read sets, we did three analyses:

1. We aligned reads in all 1,422 query-Run pairs with perfect Pebblescout score by BLAST. To assess the coverage of the query by a subject, for each pair, we used alignments >=75 bp long, or align without any overhangs except at the end of queries. We did not restrict alignments by percent identity as several read sets are from long-read nanopore sequencing with high error rates. We found alignments to the entire query in all 1,422 pairs.
2. For the 408 out of 1,422 pairs with a perfect Pebblescout score where the SRA run has long reads sequenced using nanopore, the number of reads where each read aligns to the entire length of the query varies from 1 to 158 with a median of 40 for the 408 pairs.
3. We assembled 1,014 out of 1,422 pairs with a perfect Pebblescout score where the SRA run has short reads sequenced using Illumina. The best scoring run for mecC was ERR5194629, with score of 71.24, and two runs, SRR11482358 and ERR2587962, tied for best score of 96.84 to mecD. Assembly was done using SAUTE^23^ with parameter “min_count 1” as the runs are metagenomic. Option “min_hit_len 100” was also added for the three runs that did not have perfect Pebblescout score as we do not necessarily expect full-length assemblies for those runs.

Out of 1,014 query-subject pairs with a perfect Pebblescout score for metagenomic runs with short reads, 1,007 are to mecA variants and 7 are to mecB.1. Out of the 1,007 query-subject pairs to mecA variants, 990 pairs have a full-length alignment in a single contig and 17 have alignments split on more than one contig or bases missing in the assembly. Out of 990 pairs, 555 align at 100% identity. All seven runs with a perfect Pebblescout score to mecB.1 have a contig with full length alignment to mecB.1. Six align at 100% identity, and the best matching contig in SRR11482360 assembly has two mismatches. Five reads support one mismatch, but only one read supports the second mismatch.

For mecC and mecD that did not have any runs with a perfect Pebblescout score, we find the following using assemblies generated as described above. ERR5194629 has contigs that align to 82.6% of mecC at an average alignment identity of 99%. Contigs in SRR11482358 and ERR2587962 align to 88.3% and 85.7% of mecD at an average alignment identity of 99.81% and 99.84%, respectively. These results show that Pebblescout identifies excellent candidates for use in assembling antimicrobial resistance genes. It also shows that Pebblescout finds genes that are split into contigs as the working unit for Pebblescout is kmers and not reads or contigs.

### Find subjects with SNPs of interest (*Acinetobacter baumannii* example)

Whole genome sequencing of cultured isolates is used for detecting pathogens in GenomeTrakr and NCBI’s Pathogen Detection Pipeline^34^. For pre-screening the clusters, it is desirable to find single nucleotide polymorphisms (SNPs) that can act as *cluster signatures* for distinguishing one cluster from the rest. Six SNPs from the reference genome for ACICU (accession CP000863.1) reported^35^ differentiate two published outbreak clusters of the clinically relevant bacterial pathogen *Acinetobacter baumannii* from non-cluster isolates.

Using the reference and variant bases provided by the authors in the study^35^ for each of the six SNPs and extending them by three different lengths on both ends using the reference genome sequence, we generated three sets of 12 queries. Extension by 100 bp, 41 bp, and 24 bp on both ends of the polymorphic base were used to generate *Long, Middle,* and *Short* sets of 12 queries each with length 201 bp, 83 bp, and 49 bp, respectively.

Table S9 shows the number of RefSeq assemblies, WGS assemblies, and metagenomic runs that have a perfect Pebblescout score for each query in each set without any masking of kmers. For all three sets, the same set of assemblies was reported with a perfect Pebblescout score in the search to the RefSeq database. The same set was reported in search against the WGS database for Long and Middle query sets, but the Short set reported one additional assembly (JJRW01 for query cluster2.SNP5.varied). As expected per the results reported by the authors of the study^35^, the sets of assemblies found using variant base for SNP3, SNP4, and SNP5 in cluster2 (i.e., queries cluster2.SNP3.varied, cluster2.SNP4.varied, and cluster2.SNP5.varied) are subsets of assemblies for SNP6 in cluster2. We verified by BLAST that all 41 and 74 assemblies reported using the Long queries for RefSeq and WGS, respectively, have the variant base at the SNP position in these assemblies. JJRW01 found using the short query appears to be a false positive as it only has 100% identity matches of length 26 around the SNP position.

**Table S9:** Number of RefSeq assemblies, WGS assemblies, and metagenomic Runs that have a perfect Pebblescout score for each query in each set. Meta in the table is short for metagenomic.

| SNP | Long |  |  | Middle |  |  | Short |  |  |
| --- | --- | --- | --- | --- | --- | --- | --- | --- | --- |
|  | RefSeq | WGS | Meta | RefSeq | WGS | Meta | RefSeq | WGS | Meta |
| cluster1.SNP1.ACICU | 5562 | 11305 | 907 | 5978 | 11861 | 2499 | 6005 | 11897 | 2936 |
| cluster1.SNP2.ACICU | 4132 | 8492 | 673 | 4680 | 9326 | 1103 | 4689 | 9343 | 1235 |
| cluster2.SNP3.ACICU | 3722 | 7948 | 191 | 4480 | 9399 | 1086 | 5907 | 11585 | 2638 |
| cluster2.SNP4.ACICU | 4124 | 8877 | 594 | 5195 | 10789 | 1785 | 5784 | 11671 | 2299 |
| cluster2.SNP5.ACICU | 3835 | 7984 | 185 | 4547 | 9175 | 844 | 4656 | 9315 | 1002 |
| cluster2.SNP6.ACICU | 5762 | 11553 | 991 | 5781 | 11583 | 1445 | 5805 | 11622 | 1753 |
| cluster1.SNP1.varied | 0 | 0 | 54 | 0 | 0 | 82 | 0 | 0 | 86 |
| cluster1.SNP2.varied | 0 | 0 | 33 | 0 | 0 | 67 | 0 | 0 | 67 |
| cluster2.SNP3.varied | 1 | 1 | 3 | 1 | 1 | 13 | 1 | 1 | 20 |
| cluster2.SNP4.varied | 1 | 1 | 11 | 1 | 1 | 24 | 1 | 1 | 46 |
| cluster2.SNP5.varied | 15 | 23 | 6 | 15 | 23 | 11 | 15 | 24 | 13 |
| cluster2.SNP6.varied | 41 | 74 | 8 | 41 | 74 | 9 | 41 | 74 | 22 |

The number of matches to metagenomic runs in Long, Middle, and Short queries with perfect Pebblescout score is 115, 206, and 254 coming from 105, 183, and 215 distinct runs, respectively. We verified that all 115 matches reported using Long queries are correct as there is at least one 42-mer supporting the variant base for all 115 matches. Only four out of 206 and ten out of 254 matches do not have support for the variant base and are false positives due to very few kmers sampled from the queries.

This analysis shows we can find subjects with SNPs of interest by creating queries using the SNP base and that the false positive rate is low in the matches reported with perfect Pebblescout score. We note that while having longer extensions around the SNP base reduces false positives, it also precludes finding subjects if another SNP is present in the sequences in the extension.

### Find similar large and heterogeneous samples (microbial community example)

Sequences from a recent study^36^ analyzing microbial processes occurring in low-oxygen bottom waters of the Chesapeake Bay are available in BioProject PRJNA612045. For generating the query set, SRR14874039 from PRJNA612045 was suggested by Dr. Sarah Preheim who co-authored the study^36^. Given the limit of 15 megabytes on the Pebblescout web page and 150 bp length for reads in SRR14874039, we randomly selected one hundred thousand reads from SRR14874039 and concatenated them with an ambiguity character ‘N’ to generate one query of length 15.1 megabases (14.5 megabytes text file). Searching metagenomic database with this query and categorizing location for top 50 scoring samples shows that we have 30 samples from Chesapeake Bay, 10 from Skidaway Island, Georgia, 7 from Delaware River and Bay, 2 from Pivers Island, North Carolina, and one from Black Sea, Varna Bay Bulgaria (Sample SAMEA7392479, Run ERR4674194). That is, all except ERR4674194 appear to be geographically close to Chesapeake Bay in Maryland.

In this application, a second question which can be asked is to find runs specific to the Chesapeake Bay. To do so requires giving higher weight to kmers found in few samples and downweighing, essentially ignoring, kmers seen in many samples. Changing the scoring constant to one with rest of the input as before shows that locations for top 50 scoring samples changes to 45 samples from the Chesapeake Bay, 2 from the Delaware River and Bay, and one each from Pivers Island, North Carolina, Barataria Bay, Louisiana, and Black Sea, Varna Bay Bulgaria.

These results suggest that Pebblescout identifies runs with microbial community similar to the one in the input query and allows for different questions to be answered for the same input with different parameters, when the size and properties of the input support finding such differences.

### Pilot web service and searches for applications

The web page for searching the six databases is at https://pebblescout.ncbi.nlm.nih.gov/. A user provides nucleotide query sequence that is at least 42 bases long, chooses the database, selects the type of search they wish to perform, changes parameters as appropriate, and views or downloads the output. Query sequences can be typed or pasted in the provided text box or uploaded as a FASTA format file. Queries can also be provided as a comma separated list of GenBank accessions in the text box.

To maintain the interactive nature of the website, some limits are imposed that should not affect most users but avoids abuse of the current system that supports the web page. The following limitations are currently in place:

1. The default for “Max #subjects per query” is 100, but a user can change that to a maximum value of 100,000.
2. The maximum number of rows printed is 1,000 in the “View” output and one million in the “Download” output.
3. Masking limits can be increased above the default value shown for each database. However, the maximum value allowed on the web page for a query is computed based on the number of expected results and memory required for the same. If the specified value is above the computed limit, an error message is printed, specifying the maximum allowed limit for the query. See Figure S3 for an example.
4. A user must re-click View or Download after parameters are changed.
5. The size of the query on the website is limited to 15 megabytes (15,728,640 characters) to accommodate up to 15 megabase sequences and additional characters for the definition lines.
6. A user must specify a list of subjects for Detailed search.

**Figure S3:**
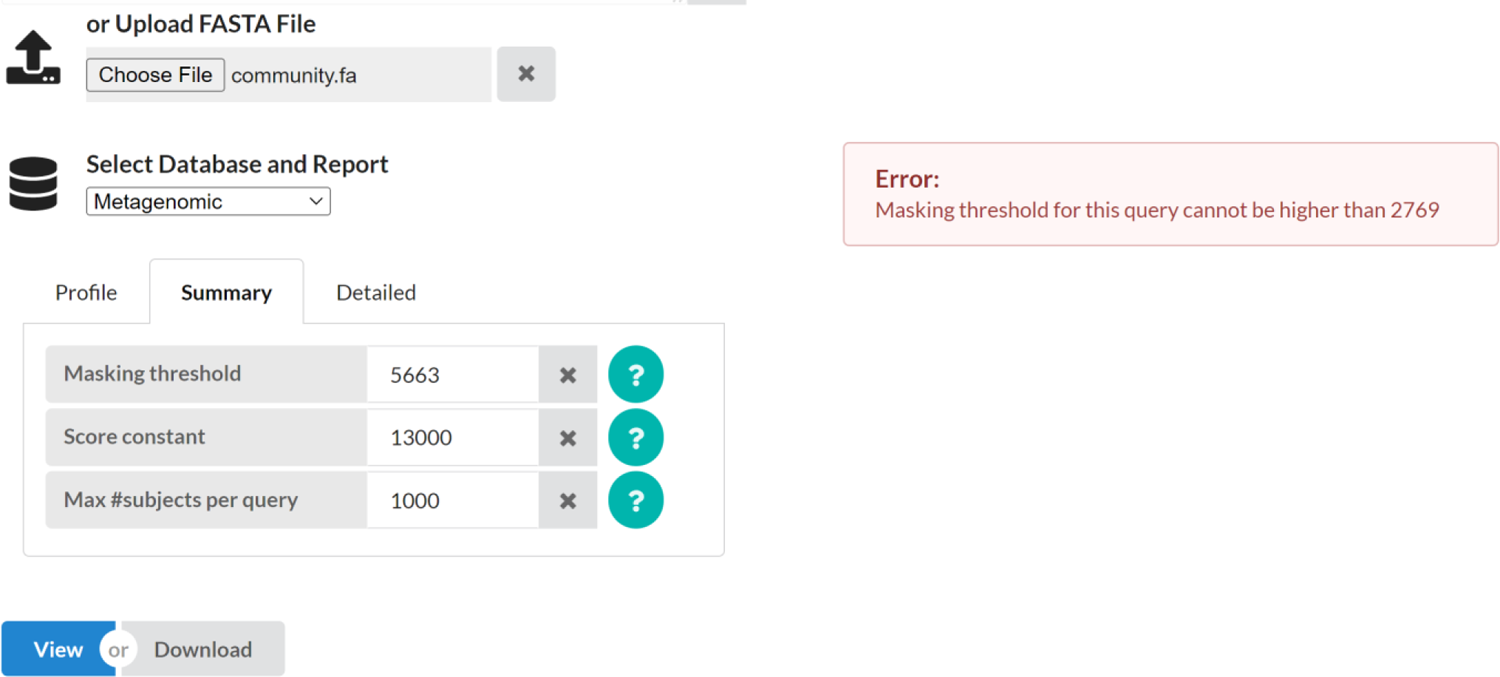
Error message with default masking threshold for application S6.

For applications presented in this manuscript, files containing input sequences for all queries and output of the searches are provided as stated in the code and data availability section. Here, we collate database and parameter settings for all applications so results reported can be reproduced using the web page (Table S10). The six research applications are numbered R1 through R6. We also provide the wall clock time for each query when we executed them on the web. It is difficult to control cache behavior for the searches, especially on the web. We spread out the searches inhouse over several days and did the same search three times with the second and third search starting right after the previous one. Wall clock time for inhouse searches show the effect of cache on the search time.

As the limit on the web page for the size of input is 15 megabytes and number of rows that can be downloaded is up to one million, we split the queries for some applications into multiple files such that each query is present in exactly one file. Doing so allows one to achieve the same results we get when analysis was done inhouse using a single input file for each application.

For application R4 and R5 where we wish to use all kmers sampled from the query, we disable masking by setting it to a number larger than the number of subjects in the database. For application R6, the maximum value allowed for masking on the web is 2769. We note that the top 50 results do not change for this application when we do the search in-house with the default of 5663. For users who need to use parameters not supported on the web page, please contact NCBI at with information about your application and justification for the choice of parameters.

**Table S10:** For all applications, the queries used, search performed, wall clock time for the search on the web, and wall clock time for three consecutive runs done inhouse are listed. Six research applications are prefixed with letter ‘R’. Accession list in application R1 for detailed search is the list of runs in Table S7: ERR2241729, ERR4083961, SRR11414099, SRR14160768, SRR16071456, SRR3176533, SRR4425666, SRR8925718, SRR12874575. For all output, Download and the maximum subject count of 100,000 was specified. Any other non-default parameters are listed.

| Query |  |  | Web search | Inhouse search wall clock time (sec) |  |  | Search |  |  |
| --- | --- | --- | --- | --- | --- | --- | --- | --- | --- |
| Fasta file | Base count | Seq count | Wall clock (sec) | 1 | 2 | 3 | Database | Mode | Parameters |
| **Application 1: Find previously undiscovered species related to known species** |  |  |  |  |  |  |  |  |  |
| serratus.1 | 4320363 | 648 | 40.15 | 154.53 | 4.82 | 4.49 | Metagenomic | Summary |  |
| serratus.2 | 4247800 | 700 | 53.95 | 136.7 | 4.97 | 4.97 | Metagenomic | Summary |  |
| serratus.3 | 4648502 | 662 | 61.98 | 138.45 | 4.78 | 4.35 | Metagenomic | Summary |  |
| serratus.4 | 4290439 | 570 | 38.86 | 111.69 | 4.48 | 4.21 | Metagenomic | Summary |  |
| serratus.5 | 3833989 | 595 | 28.43 | 106.33 | 4.34 | 5.64 | Metagenomic | Summary |  |
| serratus.6 | 4310373 | 668 | 44.12 | 128.98 | 5.18 | 4.79 | Metagenomic | Summary |  |
| serratus.7 | 3885619 | 625 | 76.62 | 95.86 | 4.07 | 4.37 | Metagenomic | Summary |  |
| serratus.8 | 4451562 | 678 | 93.34 | 124.43 | 4.64 | 4.56 | Metagenomic | Summary |  |
| serratus.9 | 3915021 | 570 | 72.21 | 88.6 | 4.59 | 4.59 | Metagenomic | Summary |  |
| serratus.10 | 4113490 | 701 | 71.49 | 88.42 | 5.63 | 4.57 | Metagenomic | Summary |  |
| serratus.11 | 4376661 | 514 | 36.76 | 128.35 | 9.63 | 9.16 | Metagenomic | Summary |  |
| serratus.12 | 3823679 | 783 | 3.59 | 91.66 | 6.46 | 7.95 | Metagenomic | Summary |  |
| serratus.13 | 3352961 | 372 | 3.52 | 94.65 | 6.97 | 6.6 | Metagenomic | Summary |  |
| serratus.14 | 3979212 | 609 | 3.70 | 100.72 | 6.85 | 8.31 | Metagenomic | Summary |  |
| serratus.15 | 4305018 | 715 | 83.23 | 98.62 | 8.02 | 5.69 | Metagenomic | Summary |  |
| serratus.16 | 4367890 | 581 | 3.68 | 118.97 | 6.67 | 6.27 | Metagenomic | Summary |  |
| serratus.17 | 3809950 | 586 | 3.28 | 91.85 | 6.95 | 6.94 | Metagenomic | Summary |  |
| serratus.18 | 3865432 | 544 | 4.61 | 92.57 | 4.72 | 4.3 | Metagenomic | Summary |  |
| serratus.19 | 4440127 | 761 | 3.60 | 97.14 | 4.47 | 5.03 | Metagenomic | Summary |  |
| serratus.20 | 2711241 | 413 | 28.50 | 47.59 | 3.09 | 2.91 | Metagenomic | Summary |  |
| **Application 2: Find known species in new environments** |  |  |  |  |  |  |  |  |  |
| candida.its | 320 | 1 | 0.11 | 0.05 | 0.08 | 0.06 | Metagenomic | Profile |  |
| candida.its | 320 | 1 | 0.09 | 0.11 | 0.05 | 0.05 | Metagenomic | Summary | Mask: 76 |
| **Application 3: Find host species for endosymbionts** |  |  |  |  |  |  |  |  |  |
| wolbachia | 30268 | 44 | 0.50 | 0.79 | 0.38 | 0.38 | Metagenomic | Summary |  |
| **Application 4: Find assemblies in WGS for additional context or to support novel sequence findings** |  |  |  |  |  |  |  |  |  |
| ssRNA.1 | 7912069 | 7845 | 2.08 | 133.86 | 2.33 | 2.41 | WGS | Summary |  |
| ssRNA.2 | 8034957 | 7766 | 4.41 | 74.79 | 2.46 | 2.44 | WGS | Summary |  |
| **Application R1: Detect hard-to-sequence samples** |  |  |  |  |  |  |  |  |  |
| cc.mrna.all | 1.1E+07 | 5793 | 5.70 | 648.46 | 5.2 | 4.8 | Metagenomic | Detailed | Accession list |
| cc.mrna.1 | 889084 | 440 | 35.36 | 40.62 | 2.47 | 3.13 | Metagenomic | Summary |  |
| cc.mrna.2 | 784494 | 386 | 2.26 | 36.41 | 2.74 | 2.55 | Metagenomic | Summary |  |
| cc.mrna.3 | 775752 | 399 | 2.11 | 35.88 | 2.8 | 2.47 | Metagenomic | Summary |  |
| cc.mrna.4 | 766289 | 414 | 2.20 | 35.27 | 3.24 | 3.1 | Metagenomic | Summary |  |
| cc.mrna.5 | 777375 | 432 | 2.23 | 35.04 | 2.89 | 2.58 | Metagenomic | Summary |  |
| cc.mrna.6 | 723638 | 389 | 2.11 | 32.55 | 3.09 | 2.63 | Metagenomic | Summary |  |
| cc.mrna.7 | 773752 | 381 | 2.26 | 36.44 | 3.65 | 2.52 | Metagenomic | Summary |  |
| cc.mrna.8 | 665105 | 368 | 2.19 | 30.89 | 2.83 | 2.27 | Metagenomic | Summary |  |
| cc.mrna.9 | 651591 | 363 | 2.13 | 29.12 | 2.64 | 2.25 | Metagenomic | Summary |  |
| cc.mrna.10 | 803419 | 448 | 2.18 | 35.25 | 2.8 | 2.45 | Metagenomic | Summary |  |
| cc.mrna.11 | 788190 | 406 | 2.26 | 34.62 | 2.78 | 2.99 | Metagenomic | Summary |  |
| cc.mrna.12 | 842171 | 425 | 2.18 | 36.82 | 3.1 | 2.78 | Metagenomic | Summary |  |
| cc.mrna.13 | 813125 | 396 | 33.01 | 34.44 | 2.45 | 2.59 | Metagenomic | Summary |  |
| cc.mrna.14 | 768558 | 384 | 2.13 | 32.62 | 2.43 | 2.74 | Metagenomic | Summary |  |
| cc.mrna.15 | 305480 | 162 | 1.02 | 13.17 | 1.06 | 0.94 | Metagenomic | Summary |  |
| **Application R2: Find variant/invariant regions of genomes** |  |  |  |  |  |  |  |  |  |
| ecoli_k12 | 4641652 | 1 | 2.24 | 54.29 | 2.16 | 2.15 | Metagenomic | Profile |  |
| HIV_1 | 9181 | 1 | 0.70 | 1.37 | 0.12 | 0.1 | Human<br>RNAseq | Profile |  |
| **Application R3: Find contamination** |  |  |  |  |  |  |  |  |  |
| cov2 | 29903 | 1 | 1.24 | 0.96 | 0.07 | 0.07 | WGS | Summary |  |
| **Application R4: Finding subjects with genes of interest** |  |  |  |  |  |  |  |  |  |
| mec | 29855 | 14 | 1.24 | 1.88 | 0.61 | 0.47 | RefSeq | Summary | Mask: 3500000 |
| mec | 29855 | 14 | 1.92 | 1.79 | 1.28 | 1.2 | WGS | Summary | Mask: 3500000 |
| mec | 29855 | 14 | 1.28 | 1.46 | 1.12 | 1.06 | Metagenomic | Summary | Mask: 3500000 |
| **Application R5: Find subjects with SNPs of interest** |  |  |  |  |  |  |  |  |  |
| snps_long | 2412 | 12 | 0.31 | 0.49 | 0.22 | 0.23 | RefSeq | Summary | Mask: 3500000 |
| snps_middle | 996 | 12 | 0.24 | 0.3 | 0.17 | 0.16 | RefSeq | Summary | Mask: 3500000 |
| snps_short | 588 | 12 | 0.17 | 0.43 | 0.1 | 0.1 | RefSeq | Summary | Mask: 3500000 |
| snps_long | 2412 | 12 | 0.56 | 0.42 | 0.41 | 0.4 | WGS | Summary | Mask: 3500000 |
| snps_middle | 996 | 12 | 0.43 | 0.29 | 0.28 | 0.28 | WGS | Summary | Mask: 3500000 |
| snps_short | 588 | 12 | 0.22 | 0.18 | 0.17 | 0.14 | WGS | Summary | Mask: 3500000 |
| snps_long | 2412 | 12 | 0.43 | 0.32 | 0.32 | 0.31 | Metagenomic | Summary | Mask: 3500000 |
| snps_middle | 996 | 12 | 0.23 | 0.16 | 0.15 | 0.15 | Metagenomic | Summary | Mask: 3500000 |
| snps_short | 588 | 12 | 0.13 | 0.08 | 0.08 | 0.11 | Metagenomic | Summary | Mask: 3500000 |
| **Application R6: Find similar large and heterogeneous samples** |  |  |  |  |  |  |  |  |  |
| community | 1.5E+07 | 1 | 7.65 | 441.8 | 8.06 | 8.04 | Metagenomic | Summary | Mask: 2769 |
| community | 1.5E+07 | 1 | 7.81 | 567.65 | 6.74 | 6.7 | Metagenomic | Summary | Mask: 2769,<br>Constant: 1 |

## Notes

https://pebblescout.ncbi.nlm.nih.gov

https://ftp.ncbi.nlm.nih.gov/pub/agarwala/pebblescout/v2.25_release.zip

### Competing Interest Statement

The authors have declared no competing interest.

